# Recovering birds and mammals across Europe continue to be negatively impacted by threats but benefit from conservation measures

**DOI:** 10.1101/2022.09.01.506271

**Authors:** Claudia Gray, Louise McRae, Stefanie Deinet, Sophie Ledger, Charlotte Benham, Ian J. Burfield, Mark Eaton, Kate Scott-Gatty, Hannah Puleston, Claire Rutherford, Anna Staneva, Frans Schepers, Robin Freeman

## Abstract

Even during ongoing global biodiversity losses and extinctions, numerous species have shown recoveries in terms of increased abundance and/or range extent. Understanding the mechanisms that contribute to, or limit, these recoveries is critical not just to ensure they continue, but to promote similar recoveries across broader ecosystems. Here, we explore the changes in abundance and range extent of selected 47 recovering species (24 mammals and 23 birds) in Europe using official data reported by EU Member States and supplemented using the Living Planet Index database. We investigate how the diversity of ongoing threats and conservation measures contribute to the likelihood and extent of recoveries. For birds, long-term recoveries were less likely among species impacted by a greater diversity of threats, although this may be mitigated by the diversity of conservation measures in place. Similarly, for mammals, populations with reported threats recovered less quickly while those with management actions in place recovered more quickly. To achieve the aims of the UN Decade on Restoration, we need to ensure, *even for recovering species*, that threats continue to be reduced and that conservation management actions are ongoing and effective.

## Introduction

It is now widely accepted that biodiversity, and therefore also ecosystem services and nature’s contributions to people, is deteriorating worldwide at an unprecedented speed (Díaz et al. 2019). Humanity has responded with action all the way from the grassroots, local level up to the establishment of, and growing collaboration between, international bodies and conventions. The Intergovernmental Science-Policy Platform on Biodiversity and Ecosystem Services (IPBES) and Intergovernmental Panel on Climate Change (IPCC) have now jointly recognised the interconnectedness of the biodiversity and climate change crises (Pörtner et al. 2021). We have recently entered the UN Decade on Ecosystem Restoration (decadeonrestoration.org), with the explicit aim “to prevent, halt, and reverse the degradation of ecosystems on every continent and in every ocean”. Although this is a great challenge, we already see multiple sources of hope. For example, the conservation optimism movement (https://conservationoptimism.org/) focuses on success stories, which can be crucial to drive engagement and action (Mcafee et al. 2019). Similarly, the concept of rewilding has provided a vision for how even degraded landscapes can regain key ecological functions and reinvigorate the connections between people and nature (Schepers & Jepson 2016); this has been particularly powerful in Europe where we have few, if any, truly wild habitats left.

If we are to continue to improve efforts to drive the recovery of biodiversity worldwide, the ecosystems of Europe provide a particularly interesting area on which to focus. The continent has experienced high levels of habitat conversion and is therefore an extreme case in terms of the extent of the recovery required. Large-scale habitat destruction, agricultural expansion, widespread hunting of ungulates and persecution of large carnivores prior to the 20^th^ Century contributed to the loss and near extinction of many native European species (Chapron et al. 2014; Crees et al, 2019; Linnell et al. 2020). Similar factors have driven declines in many bird species across Europe, with agricultural intensification having a widespread negative impact on farmland bird species in the later part of the 20^th^ Century (Donald et al. 2001; Bowler et al. 2021).

However, we are now seeing the recovery of numerous species of birds, mammals and other taxonomic groups (e.g. Deinet et al. 2013; Tucker et al. 2019). These recoveries are the result of several factors. In response to a variety of growing pressures, the European Union (EU) established the Nature Directives, the Birds Directive (1979) and Habitats Directive (1992). These form the primary legislation protecting biodiversity in Europe and contributing to the EU’s implementation of the Bern Convention and the Convention on Biological Diversity (Epstein et al. 2016). The Nature Directives aim to restore habitats and species to “Favourable Conservation Status” and established Natura 2000, the network of European protected areas. Alongside these key legislative changes, Europe has also experienced widespread rural-urban migration, agricultural abandonment, forest regeneration, active ecosystem restoration and purposeful reintroductions.

To ensure the ongoing recovery of European biodiversity, it is essential that we continue to improve our understanding of the factors facilitating or limiting species recoveries. Populations of species across Europe will face a variety of threats and conservation measures, and understanding how these interact to contribute to or curtail recoveries is critical. Here, we use a combination of data reported under the EU Nature Directives and the Living Planet Database (LPD) to explore these factors and how they influence European wildlife recovery.

The EU Nature Directives coordinate conservation efforts for more than 2,000 species and habitats across the EU Member States, with the aim of maintaining or restoring a favourable conservation status (European Environment Agency 2020a). Member States are required to report every six years on the sizes of and trends in populations of birds (under Article 12 of the Birds Directive) and on the conservation status of and trends in targeted habitats and species (under Article 17 of the Habitats Directive). The European Environment Agency (EEA) and its European Topic Centre on Biological Diversity, alongside several other working groups, provide technical support to Member States and review the data reported. Reports from Member States now provide a baseline for measuring the status of and trends in species and habitats over multiple monitoring periods. Under the Habitats Directive, species data have been collected for three reporting periods (2000 - 2006, 2007 - 2012, and 2013 - 2018). Under the Birds Directive, data have been collected for two reporting periods (2008 - 2012 and 2013 - 2018). These data underpin the State of Nature in the EU report (European Environment Agency 2020a) and are used to inform EU policies, measure progress towards EU strategy targets and facilitate monitoring of the EU’s contribution to the Sustainable Development Goals and Convention on Biological Diversity. For birds, the same data are combined with analogous data from non-EU countries to assess the conservation status of species at EU and European scales in the European Red List of Birds (BirdLife International, 2021).

For less systematically monitored species, trends in population abundance may only be available from specific sources – e.g. research articles, governments or NGO reports. The Living Planet Database (LPD) collates many of these sources into a single framework, providing aggregated trends in abundance for over 5,000 vertebrate species globally. The individual time series of vertebrate population sizes (or proxies) from around the world are used to calculate the Living Planet Index (LPI) (Loh et al. 2005; Collen et al. 2009; McRae et al. 2017). These data underpin a variety of assessments of how threats influence wildlife population trends (Spooner et al, 2018; Daskalova et al, 2020; Green et al, 2020; Capdevilla et al, 2022; Williams et al, 2022), on the effectiveness of conservation management in protected areas (Craigie et al. 2010, Jellesmark, 2021) and to test policy scenarios (Nicholson et al. 2012; Leclère et al. 2020)

Using these available data on abundance and range, we assessed the extent to which conservation measures and threats influence the trajectories of specific bird and mammal species currently recovering in Europe. Here, we present the results from three sets of analyses on three different sets of data. First, we used the data reported by EU Member States under Article 12 of the Birds Directive to test for a relationship between the diversity of threats and conservation measures and the change in population and range size of selected breeding bird species. Second, we used the data reported under Article 17 of the Habitats Directive to test for a relationship between the diversity of threats and conservation measures and the change in population and range size of selected mammal species. Third, we augmented the analysis of recovering mammal species by using the Living Planet Database to test for a relationship between population level abundance and key variables capturing threats and conservation measures.

## Methods

### Species selection

We were specifically interested in species that are already considered to be recovering within Europe. We therefore used the species list from the 2013 “Wildlife Comeback in Europe” report and explored whether any additional species could now be added using the criteria that they had “all undergone a recovery after a period of serious decline” (Deinet et al. 2013). For birds, this allowed us to add another five species, based on information in the latest European Red List (BirdLife International 2021) and European Breeding Bird Atlas (Keller et al. 2020). For mammal species, we used the LPI data, as this was the most detailed, to confirm populations were increasing overall across a continuous data coverage of at least 2 years and up to 64 years. This process gave us a set of 23 bird and 24 mammal focal recovering species as detailed in Table S1.

### Bird data from Article 12

Article 12 data for birds reported by EU Member States in 2019 were downloaded from www.eea.europea.eu on 01/04/2021 (European Environment Agency 2021a). The dataset includes information on trends in population size and range (both long and short term), current threats (factors currently affecting the species) and future threats (factors that will affect the species in future), and conservation measures per country. Note that within the Article 12 data current threats are referred to as *pressures* and future threats as *threats*. For simplicity and alignment with the mammal population data used later in this analysis, we use the term *threats* throughout, but are referring to current ongoing pressures/threats.

We extracted breeding season data on population and range change, threats and measures reported for the 23 focal species (see Table S1) only. These data included 32 countries and territories: all current EU Member States, plus the UK, and Gibraltar, the Azores, Madeira, and the Canary Islands (reporting for Article 12 considers these separately to the UK, Portugal and Spain respectively). In these data, a population represents the data reported by the EU Member State for the entire breeding population in the country, and between one and 28 countries reported data for each species (see Table S1).

For each species within each country, we calculated the diversity of threats and conservation measures listed, as the sum of the number of level 1 pressure categories listed (i.e. the broader category groups, Table S4). We decided against using the count of the finer level (level 2) categories, as this would not capture the diversity of threats or measures (a given count could result from a set of similar threats/measures from within the same level 1 category (e.g A01, A02, A03) or from several different level 1 categories, e.g A, B, C). We removed codes starting with X (Unknown pressures, no pressures and pressures from outside the Member State) and removed measure codes starting with CX (measures outside the Member State) to avoid double counting of measures across Member States.

We calculated four response variables. The first was a binary variable capturing whether the population size was reported as increasing or decreasing, henceforth referred to as P(increasing). The second response variable was the magnitude of change in population size. Where two values for minimum and maximum magnitude of change were given, we used the mean of these values. The third was a binary variable capturing whether the species’ range was reported as expanding or shrinking, henceforth referred to as P(expanding). The fourth was the magnitude of change in the range of the species (as before, we used the mean of the maximum and minimum value where these were reported separately). In all cases, we excluded populations with statuses of fluctuating and stable so that the meaning of the binary variables P(increasing) and P(expanding) could be clearly interpreted, and so that positive or negative values for magnitude change could be easily interpreted (positive and negative magnitude change values for fluctuating or stable populations may not necessarily be interpreted in the same way). For all response variables, data were reported for a short-term time span (approx. 2007 to 2018, with an average span of 11 years, although there is some variation around these dates, see Fig S1) and a long-term time span (approx.. 1980 to 2018, with an average span of 35.1 years, although again there is considerable variation around these dates, see Fig S1).

We used linear mixed effects models to test for a relationship between each of the response variables and the number of level 1 threats and measures categories, as well as the interaction between them. We also included the length of the time span across which data were reported (number of years), to account for the considerable variation in start year and end year. The binomial family was specified for the binary response variables of P(increasing) and P(expanding). Response variables for the magnitude of population increase (Mag(increase)) and magnitude of range expansion (Mag(expansion)) were log transformed for analyses. Models were run separately for both short- and long-term data (i.e. eight analyses in total). Table S2 shows the size of datasets used for analyses of changes in selected bird species. For all models we specified random factors for the species’ identity and country. Optimal models were selected based on AIC values.

To check that results were not driven by smaller populations, which could be peripheral and less significant for the long-term recovery of the species, we repeated all 8 analyses using either weights for population size (the log transformed mean of the max and min population size variables reported) or breeding distribution surface area (also log transformed for analyses) as appropriate.

### Mammal data from Article 17

Article 17 data reported by EU Member States in 2019 (for the period 2013-2018) were downloaded from www.eea.europa.eu on 9/06/2021 (European Environment Agency 2021b). As for birds, the dataset includes information on trends in population size and range (both long and short term), threats, and conservation measures (see above for definitions of these variables).

We extracted data on the 24 focal mammal species, five of which were not included in the Article 17 data (see table S1). Depending on the species, data were available for between 2 and 54 populations, across 27 countries (all current EU Member States except Malta, plus data from the UK). In these data a population can refer to a subset of the individuals within each country, where there is more than one biogeographical or marine region in that country. For example, there are four populations of *Canis lupus* listed within France, corresponding to the regions “Atlantic”, “Alpine”, “Continental” and “Mediterranean”. There are 14 different biogeographical regions considered in these data (see (European Environment Agency 2020b) for further details).

As for birds, we summed the number of level 1 categories reported to give a diversity metric for measures and threats for each species in each country. We also removed pressure codes starting with X (Unknown pressures, no pressures and pressures from outside the Member State) and removed measure codes for species outside the Member State (CX) to avoid double counting. We calculated the same four response variables as for birds, across both the short-term time span (approx. 2000 to 2018, average span 10.4 years, see Fig S1 for details) and the long-term time span (approx. 1985 to 2018, average span 24.4 years, see Fig S1 for details).

As for birds, we used linear mixed effects models to test for a relationship between each of the four response variables, the diversity of measures and threats categories and their interaction, for both the short- and long-term time periods (eight analyses in total). Time span was also included in all models to account for variation in the number of years for which data were reported. The binomial family was specified for the binary response variables of P(increasing) and P(expanding) and the response variables of Mag(increase) and Mage(expansion) were also log transformed. Table S2 shows the size of datasets used for analyses of changes in selected mammal species. Species ID was included as a random effect in all models, but it was not possible to also include country as a random effect, or carry out sensitivity analyses in which data were weighted by population size/range size, due to the smaller size of the dataset. Model selection was based on AIC values.

### Mammal data from the LPI

To further examine the factors affecting the recovery of European mammals, we used data from the Living Planet Database (LPD), the source of species abundance data for the Living Planet Index (LPI; (WWF & ZSL 2022)). The LPD now contains over 30,000 population time series from more than 5,000 species globally. Time series information for vertebrate species is collated from published scientific literature, online databases and grey literature (government/NGO reports), totalling 4,039 individual data sources (as of Jan 2022). In these data, the definition of population for the LPI data is different from that in the EU data; a population is a species monitored at a particular location and does not always equate to the ecological definition of a population. The scale at which a population under this definition is monitored varies from site level (under a fifth of populations in the LPD) to regional (over half of populations) and national level (under a fifth of populations). Data are only included in the LPD if a measure of population size is available for at least two years, and information available on how the data were collected, what the units of measurement were, and the geographic location of the population. The data must be collected using the same method on the same population throughout the time series and the data source referenced and traceable (see (Collen et al. 2009) for further details).

We extracted data on the 24 focal mammal species (Table S1) for 1960 to 2016. Unfortunately, data were too sparse to extend the time series length to 2018 to match the Article 17 data; after 2016 these species only had data for 0 or 1 populations. Data on two populations were excluded due to creating extreme artifacts in the data that could not be verified by checking the original data sources, although there was still sufficient other population data to retain these species in the analysis. The final dataset included 940 populations in total, with between 2 and 58 populations for each species, across 38 European countries (all current EU Member States except Malta and Cyprus, plus data from the Åland Islands, Albania, Andorra, Belarus, Bosnia and Herzegovina, North Macedonia, Iceland, Norway, Serbia, Switzerland, the Russian Federation, Ukraine and the United Kingdom). Fig S2 shows the number of populations and species within the LPD for each year between 1960 and 2016. Body mass data for mammals was obtained from PanTHERIA (Jones et al. 2009).

For each population, we calculated a response variable measuring the relative change in population size since 1960: Total Lambda. Annual rates of population change are calculated following the Generalised Additive Modelling framework in Collen et al 2009, using the rlpi package (https://github.com/Zoological-Society-of-London/rlpi). The package generates a matrix of annual rates of change for each population. These annual rates were summed to give a logged value of total change in abundance for each population (Total Lambda). We calculated the length of the time series available for each population. We also obtained the following explanatory variables from the LPD, drawn originally from information provided in the source publications (further details on each are given in Table S3): a) whether the population receives targeted actions and is therefore “Managed” (Yes, No, Unknown) b) whether individuals or parts of individuals are regularly intentionally removed from the population in a way that may or may not be sustainable and/or legal, indicating that it is “Utilised” (Yes, No, Unknown), c) whether the source literature states one or more threats are known to impact the population, indicating that the population is threatened (Yes, No, Unknown), d) and whether the population is found within a protected area (Yes, No, Unknown, Both [where population spans both a protected and unprotected area]). The “Managed”, “Utilised” and “Threatened” variables do not have the option for populations to be ‘both’ as it is harder to distinguish from source publications whether these human actions only influence part of the population. Therefore, in these cases if any of the population is “Managed”, “Utilised” and “Threatened”, the variable is entered as ‘yes’.

The definition of whether a population is “Managed” within the Living Planet Database follows the relevant IUCN-CMP conservation actions and research actions classification schemes as discussed in Jellesmark et al (2022). Fig S3 shows the frequency of management actions and different threat types encoded in the LPI for these species populations.

We ran a linear mixed effects model to test for a relationship between Total Lambda and the six explanatory variables (Managed, Utilised, Threatened, Protection, Time series length and Log transformed body mass) with both species ID and Country included as random effects (n = 940 individual populations across 38 countries). Model selection was performed based on AIC values.

## Results

The threats most frequently reported under Article 12 as affecting the selected bird species were (in decreasing order): agriculture, urbanisation, exploitation, energy production and modification of water regimes (Fig 1A). These were broadly similar to the threats most commonly reported under Article 17 as affecting the selected mammal species: modification of water regimes, urbanisation, forestry, transport and exploitation (Fig 1 B). The conservation measures most commonly implemented for these birds related to these threats, although many more countries reported measures relating to mitigating pollution as a conservation measure than reported it as a pressure (Fig 1 C). Similarly, for the selected mammal species, the conservation measures reported generally corresponded to the most commonly reported threats (Fig 1 D).

**Fig 1.**
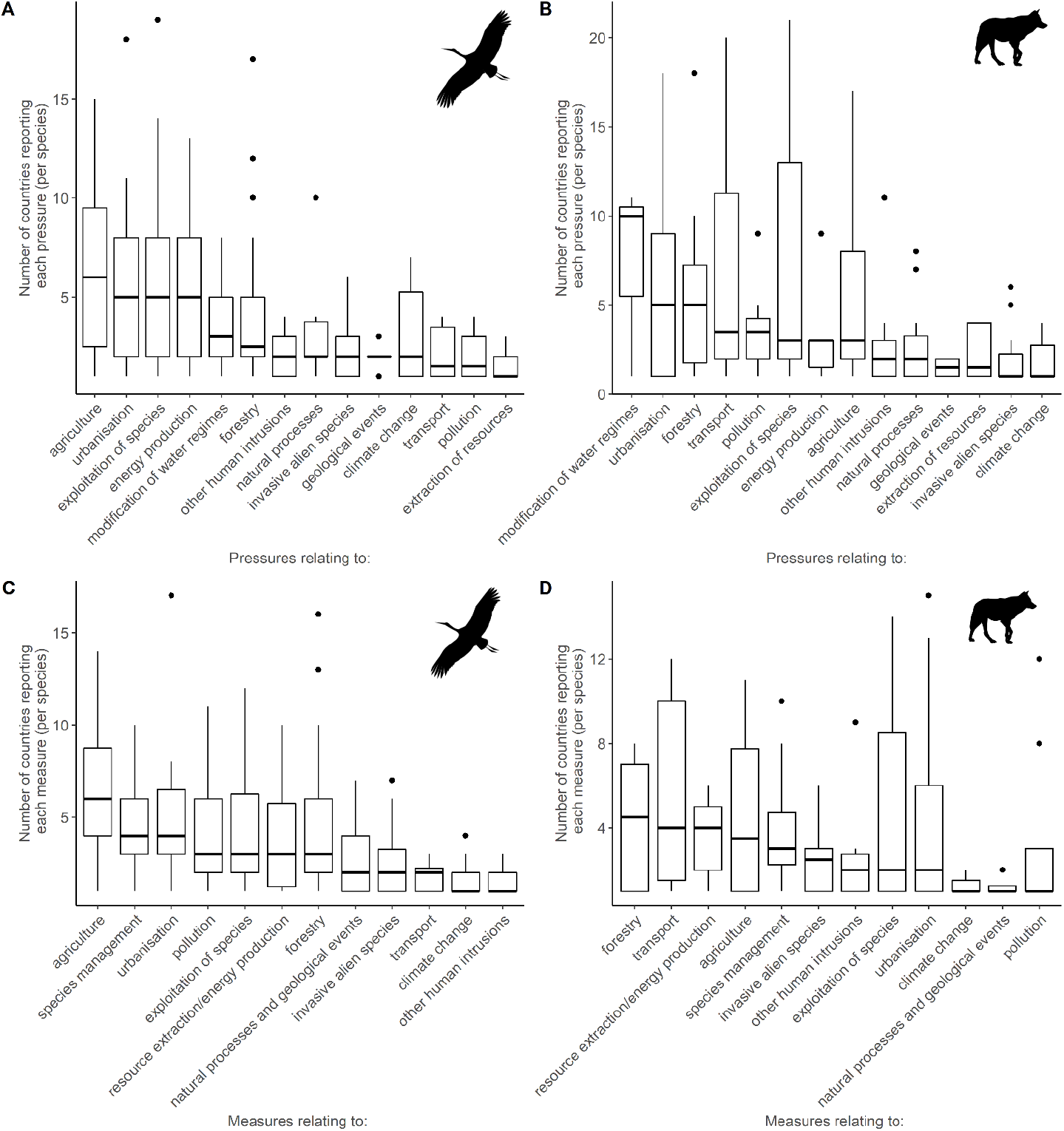
Panels show the number of countries reporting any given measure for our selected recovering bird species (A) or mammal species (B) and the number of countries reporting the corresponding threats for the selected bird species (C) and mammal species (D).

### Birds Article 12

Over the long term, there was strong evidence that as a greater diversity of threats are reported, there is a decline in the probability that bird populations increase rather than decrease (Table 1, Fig 2A). There was some evidence that this relationship may vary with the diversity of conservation measures in place (Fig S4A). We note that as the reporting period extends beyond than 40 years, P(increasing) starts to decline, but there is much higher uncertainty around the relationship due to fewer datapoints with these longer time-spans (Fig 2B and see also Fig S1A, which shows very few long-term datapoints with a start year before 1980, most long-term data is <40 years). Over the short term, however, there was no evidence for a relationship between the probability that breeding bird populations were reported as increasing (P(increasing)) and any explanatory variables (Table 1). There was no strong evidence for a relationship between the magnitude of the change in population size reported and any of the explanatory variables over the short or long term (Table 1).

**Table 1.** Model comparison for reported change in population size of selected bird species; P(increasing) = probability that the population size of the species was reported to be increasing, Mag(increase) = Magnitude of change in population size reported. Where included, the weight “log10_popsize” refers to the log transformed mean of the minimum and maximum population size reported in the Article 12 data. The best model (lowest AIC) is given in bold, multiple models are highlighted where the difference in AIC was <2. We followed a conservative approach and selected the most parsimonious or simplest model of those with deltaAIC <2.

| Response | Time Period | Formula | Weights | AIC | deltaAIC | Relative LL | AICweights |
| --- | --- | --- | --- | --- | --- | --- | --- |
| P(increasing) | short | **time_span + (1 \| speciesname) + (1 \| country)** | **none** | **114.62** | **0.00** | **1.00** | **0.48** |
| P(increasing) | short | threats + time_span + (1 \| speciesname) + (1 \| country) | none | 116.56 | 1.94 | 0.38 | 0.18 |
| P(increasing) | short | measures + time_span + (1 \| speciesname) + (1 \| country) | none | 116.61 | 1.99 | 0.37 | 0.18 |
| P(increasing) | short | measures * threats + time_span + (1 \| speciesname) + (1 \| country) | none | 117.83 | 3.21 | 0.20 | 0.10 |
| P(increasing) | short | measures + threats + time_span + (1 \| speciesname) + (1 \| country) | none | 118.56 | 3.94 | 0.14 | 0.07 |
| P(increasing) | short | **time_span + (1 \| speciesname) + (1 \| country)** | **log10_popsiz e** | **156.75** | **0.00** | **1.00** | **0.46** |
| P(increasing) | short | measures + time_span + (1 \| speciesname) + (1 \| country) | log10_popsiz e | 157.38 | 0.62 | 0.73 | 0.34 |
| P(increasing) | short | threats + time_span + (1 \| speciesname) + (1 \| country) | log10_popsiz e | 158.40 | 1.64 | 0.44 | 0.20 |
| P(increasing) | short | measures + threats + time_span + (1 \| speciesname) + (1 \| country) | log10_popsiz e | Did not converge | NA | NA | NA |
| P(increasing) | short | measures * threats + time_span + (1 \| speciesname) + (1 \| country) | log10_popsiz e | Did not converge | NA | NA | NA |
| P(increasing) | long | **threats + time_span + (1 \| speciesname) + (1 \| country)** | **none** | **114.26** | **0.00** | **1.00** | **0.31** |
| P(increasing) | long | measures * threats + time_span + (1 \| speciesname) + (1 \| country) | none | 114.26 | 0.00 | 1.00 | 0.31 |
| P(increasing) | long | measures + time_span + (1 \| speciesname) + (1 \| country) | none | 115.68 | 1.42 | 0.49 | 0.15 |
| P(increasing) | long | measures + threats + time_span + (1 \| speciesname) + (1 \| country) | none | 116.06 | 1.80 | 0.41 | 0.13 |
| P(increasing) | long | time_span_long + (1 \| speciesname) + (1 \| country) | none | 116.66 | 2.40 | 0.30 | 0.09 |
| P(increasing) | long | **threats + time_span + (1 \| speciesname) + (1 \| country)** | **log10_popsiz e** | **169.60** | **0.00** | **1.00** | **0.40** |
| P(increasing) | long | measures * threats + time_span + (1 \| speciesname) + (1 \| country) | log10_popsiz e | 169.63 | 0.03 | 0.98 | 0.39 |
| P(increasing) | long | measures + threats + time_span + (1 \| speciesname) + (1 \| country) | log10_popsiz e | 171.36 | 1.76 | 0.42 | 0.17 |
| P(increasing) | long | time_span_long + (1 \| speciesname) + (1 \| country) | log10_popsiz e | 175.29 | 5.69 | 0.06 | 0.02 |
| P(increasing) | long | measures + time_span + (1 \| speciesname) + (1 \| country) | log10_popsiz e | 176.03 | 6.43 | 0.04 | 0.02 |
| Mag(increase) | short | measures * threats + time_span + (1 \| speciesname) + (1 \| country) | none | 419.19 | 0.00 | 1.00 | 0.39 |
| Mag(increase) | short | measures + threats + time_span + (1 \| speciesname) + (1 \| country) | none | 419.43 | 0.24 | 0.89 | 0.34 |
| Mag(increase) | short | **time_span + (1 \| speciesname) + (1 \| country)** | **none** | **420.99** | **1.80** | **0.41** | **0.16** |
| Mag(increase) | short | measures + time_span + (1 \| speciesname) + (1 \| country) | none | 422.90 | 3.71 | 0.16 | 0.06 |
| Mag(increase) | short | threats + time_span + (1 \| speciesname) + (1 \| country) | none | 423.11 | 3.92 | 0.14 | 0.05 |
| Mag(increase) | short | **time_span + (1 \| speciesname) + (1 \| country)** | **log10_popsiz** | **411.12** | **0.00** | **1.00** | **0.56** |
| Mag(increase) | short | measures + time_span + (1 \| speciesname) + (1 \| country) | log10_popsiz | 413.10 | 1.98 | 0.37 | 0.21 |
| Mag(increase) | short | threats + time_span + (1 \| speciesname) + (1 \| country) | log10_popsiz | 413.15 | 2.03 | 0.36 | 0.20 |
| Mag(increase) | short | measures + threats + time_span + (1 \| speciesname) + (1 \| country) | log10_popsiz | 417.04 | 5.92 | 0.05 | 0.03 |
| Mag(increase) | short | measures * threats + time_span + (1 \| speciesname) + (1 \| country) | log10_popsiz | 420.75 | 9.63 | 0.01 | 0.00 |
| Mag(increase) | long | **time_span + (1 \| speciesname) + (1 \| country)** | **none** | **433.00** | **0.00** | **1.00** | **0.36** |
| Mag(increase) | long | measures + time_span + (1 \| speciesname) + (1 \| country) | none | 433.65 | 0.65 | 0.72 | 0.26 |
| Mag(increase) | long | threats + time_span + (1 \| speciesname) + (1 \| country) | none | 434.66 | 1.66 | 0.44 | 0.16 |
| Mag(increase) | long | measures * threats + time_span + (1 \| speciesname) + (1 \| country) | none | 434.97 | 1.97 | 0.37 | 0.13 |
| Mag(increase) | long | measures + threats + time_span + (1 \| speciesname) + (1 \| country) | none | 435.61 | 2.61 | 0.27 | 0.10 |
| Mag(increase) | long | **time_span + (1 \| speciesname) + (1 \| country)** | **log10_popsiz** | **418.88** | **0.00** | **1.00** | **0.34** |
| Mag(increase) | long | measures + time_span + (1 \| speciesname) + (1 \| country) | log10_popsiz | 419.19 | 0.31 | 0.85 | 0.29 |
| Mag(increase) | long | threats + time_span + (1 \| speciesname) + (1 \| country) | log10_popsiz | 420.78 | 1.90 | 0.39 | 0.13 |
| Mag(increase) | long | measures + threats + time_span + (1 \| speciesname) + (1 \| country) | log10_popsiz | 420.79 | 1.91 | 0.38 | 0.13 |
| Mag(increase) | long | measures * threats + time_span + (1 \| speciesname) + (1 \| country) | log10_popsiz | 421.14 | 2.26 | 0.32 | 0.11 |

**Fig 2.**
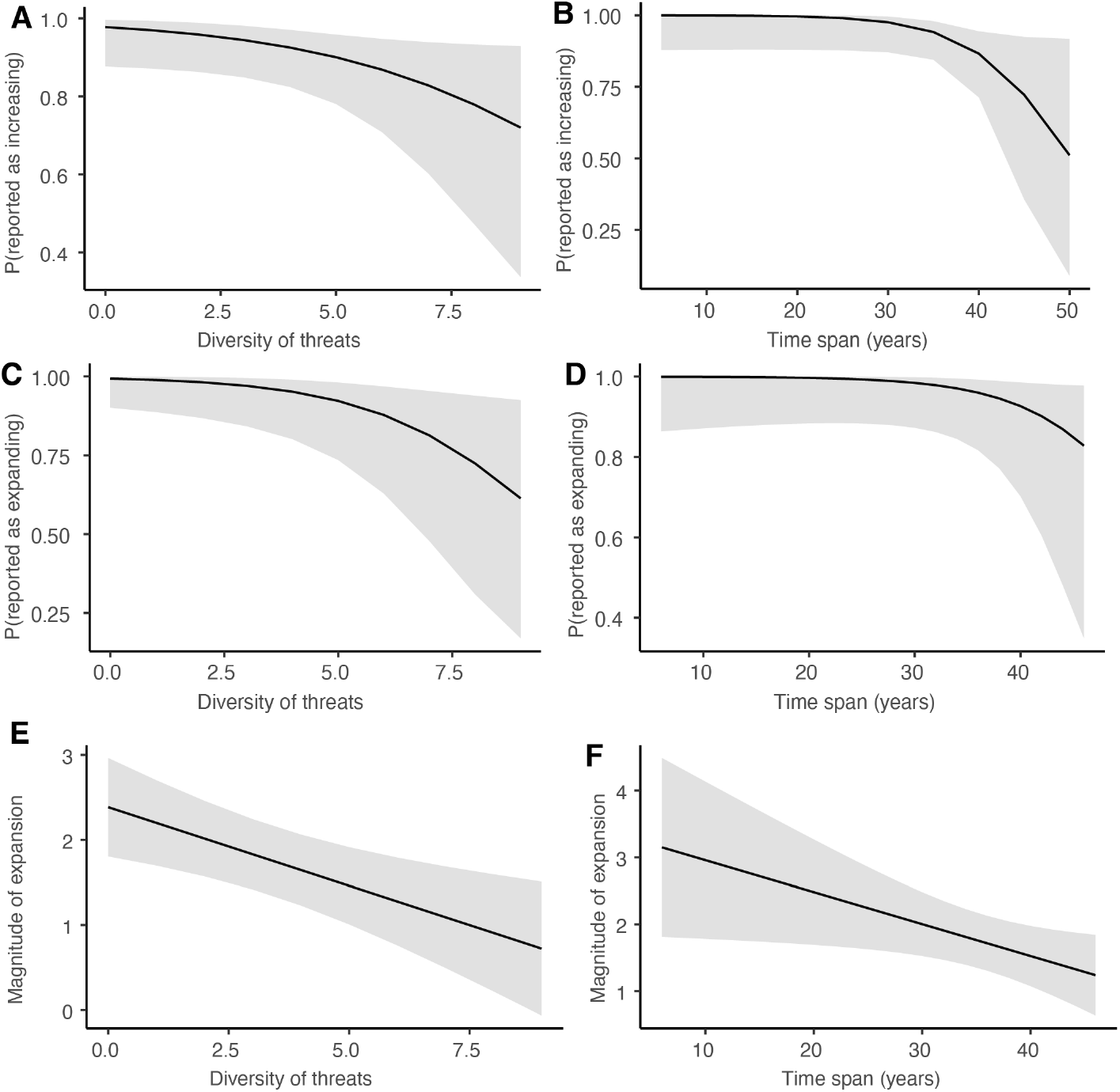
The probability that the selected breeding bird populations were reported as increasing (over the long term) gets lower as A) diversity of threats increases and B) the time span of the data increases over 40 years, but with a considerable increase in uncertainty past that 40-year span. The probability that the selected breeding bird ranges are expanding (over the long term) shows C) a decrease with a greater diversity of threats reported and D) an increase in uncertainty for time spans over 40 years. The magnitude of expansion of these bird populations also E) decreases as the diversity of threats reported increases and F) as the time span of the data increases.

Our analysis of the probability that selected bird ranges were reported as expanding was not conclusive for the short time-span, due to lack of convergence in the models run, both with and without weights for population size (Table 2, see also FigS5 A and B). However, over the long term we again found strong evidence that the probability of birds’ ranges being reported as expanding declines as a greater diversity of threats are reported (Table 2, Fig 2 C), and some evidence that this relationship depends on the diversity of conservation measures in place (Table 2, Fig S4 C). There was also an increase in uncertainty around the relationship with time span above 40 years (Fig 2 D).

**Table 2.** Model comparison for reported change in range of focal bird species; P(expanding) = probability that range of the species was reported to be expanding, Mag(expansion) = Magnitude of change in range reported. The weight “log10_surface” corresponds to the log transformed variable for breeding distribution surface area given in the Article 12 data. Best model (lowest AIC) given in bold, multiple models are highlighted where the difference in AIC was <2. If the set of best models included the null model, we follow a conservative approach and consider the null model as best.

| Response | Time Period | Formula | Weights | AIC | deltaAIC | Relative LL | AICweights |
| --- | --- | --- | --- | --- | --- | --- | --- |
| P(expanding) | short | time_span + (1 \| speciesname) + (1 \| country) | none | 113.46 | 0.00 | 1.00 | 1.00 |
| P(expanding) | short | measures + time_span + (1 \| speciesname) + (1 \| country) | none | Did not converge | NA | NA | NA |
| P(expanding) | short | threats + time_span + (1 \| speciesname) + (1 \| country) | none | Did not converge | NA | NA | NA |
| P(expanding) | short | measures + threats + time_span + (1 \| speciesname) + (1 \| country) | none | Did not converge | NA | NA | NA |
| P(expanding) | short | measures * threats + time_span + (1 \| speciesname) + (1 \| country) | none | Did not converge | NA | NA | NA |
| P(expanding) | short | measures + threats + time_span + (1 \| speciesname) + (1 \| country) | log10_surface | 202.79 | 0.00 | 1.00 | 0.58 |
| P(expanding) | short | **threats + time_span + (1 \| speciesname) + (1 \| country)** | **log10_surface** | **203.40** | **0.61** | **0.74** | **0.42** |
| P(expanding) | short | time_span + (1 \| speciesname) + (1 \| country) | log10_surface | 229.77 | 26.98 | 0.00 | 0.00 |
| P(expanding) | short | measures + time_span + (1 \| speciesname) + (1 \| country) | log10_surface | 231.74 | 28.95 | 0.00 | 0.00 |
| P(expanding) | short | measures * threats + time_span + (1 \| speciesname) + (1 \| country) | log10_surface | Did not converge | NA | NA | NA |
| P(expanding) | long | measures * threats + time_span + (1 \| speciesname) + (1 \| country) | none | 104.35 | 0.00 | 1.00 | 0.40 |
| P(expanding) | long | **threats + time_span + (1 \| speciesname) + (1 \| country)** | **none** | **104.50** | **0.16** | **0.92** | **0.37** |
| P(expanding) | long | measures + threats + time_span + (1 \| speciesname) + (1 \| country) | none | 106.22 | 1.87 | 0.39 | 0.16 |
| P(expanding) | long | measures + time_span + (1 \| speciesname) + (1 \| country) | none | 108.41 | 4.07 | 0.13 | 0.05 |
| P(expanding) | long | time_span + (1 \| speciesname) + (1 \| country) | none | 110.48 | 6.14 | 0.05 | 0.02 |
| P(expanding) | long | **threats + time_span + (1 \| speciesname) + (1 \| country)** | **log10_surface** | **103.38** | **0.00** | **1.00** | **0.53** |
| P(expanding) | long | measures + threats + time_span + (1 \| speciesname) + (1 \| country) | log10_surface | 103.63 | 0.25 | 0.88 | 0.47 |
| P(expanding) | long | measures + time_span + (1 \| speciesname) + (1 \| country) | log10_surface | 204.43 | 101.05 | 0.00 | 0.00 |
| P(expanding) | long | time_span + (1 \| speciesname) + (1 \| country) | log10_surface | 216.72 | 113.34 | 0.00 | 0.00 |
| P(expanding) | long | measures * threats + time_span + (1 \| speciesname) + (1 \| country) | log10_surface | Did not converge | NA | NA | NA |
| Mag(expansion) | short | measures * threats + time_span + (1 \| speciesname) + (1 \| country) | none | 544.31 | 0.00 | 1.00 | 0.40 |
| Mag(expansion) | short | threats + time_span + (1 \| speciesname) + (1 \| country) | none | 545.35 | 1.05 | 0.59 | 0.24 |
| Mag(expansion) | short | **time_span + (1 \| speciesname) + (1 \| country)** | **none** | **545.91** | **1.60** | **0.45** | **0.18** |
| Mag(expansion) | short | measures + threats + time_span + (1 \| speciesname) + (1 \| country) | none | 546.87 | 2.56 | 0.28 | 0.11 |
| Mag(expansion) | short | measures + time_span + (1 \| speciesname) + (1 \| country) | none | 547.82 | 3.51 | 0.17 | 0.07 |
| Mag(expansion) | short | time_span + (1 \| speciesname) + (1 \| country) | log10_surface | Inf | NA | NA | NA |
| Mag(expansion) | short | measures + time_span + (1 \| speciesname) + (1 \| country) | log10_surface | Inf | NA | NA | NA |
| Mag(expansion) | short | threats + time_span + (1 \| speciesname) + (1 \| country) | log10_surface | Inf | NA | NA | NA |
| Mag(expansion) | short | measures + threats + time_span + (1 \| speciesname) + (1 \| country) | log10_surface | Inf | NA | NA | NA |
| Mag(expansion) | short | measures * threats + time_span + (1 \| speciesname) + (1 \| country) | log10_surface | Inf | NA | NA | NA |
| Mag(expansion) | long | measures * threats + time_span + (1 \| speciesname) + (1 \| country) | none | 594.18 | 0.00 | 1.00 | 0.41 |
| Mag(expansion) | long | measures + threats + time_span + (1 \| speciesname) + (1 \| country) | none | 594.85 | 0.67 | 0.72 | 0.29 |
| Mag(expansion) | long | **threats + time_span + (1 \| speciesname) + (1 \| country)** | **none** | **596.15** | **1.97** | **0.37** | **0.15** |
| Mag(expansion) | long | measures + time_span + (1 \| speciesname) + (1 \| country) | none | 596.35 | 2.17 | 0.34 | 0.14 |
| Mag(expansion) | long | time_span + (1 \| speciesname) + (1 \| country) | none | 602.94 | 8.76 | 0.01 | 0.01 |
| Mag(expansion) | long | time_span + (1 \| speciesname) + (1 \| country) | log10_surface | Inf | NA | NA | NA |
| Mag(expansion) | long | measures + time_span + (1 \| speciesname) + (1 \| country) | log10_surface | Inf | NA | NA | NA |
| Mag(expansion) | long | threats + time_span + (1 \| speciesname) + (1 \| country) | log10_surface | Inf | NA | NA | NA |
| Mag(expansion) | long | measures + threats + time_span + (1 \| speciesname) + (1 \| country) | log10_surface | Inf | NA | NA | NA |
| Mag(expansion) | long | measures * threats + time_span + (1 \| speciesname) + (1 \| country) | log10_surface | Inf | NA | NA | NA |

There was no evidence for a strong relationship between any explanatory variables and the magnitude of the change in breeding bird range size reported over the short term (Table 2). In contrast, over the long term we found that the magnitude of the reported change in range size decreased when a greater diversity of threats was reported (Fig 2 E) and over longer time-spans (Fig 2 F). As before, there was also some evidence that the relationship with diversity of threats depends on the diversity of conservation measures (Fig S4 E).

All the results above were similar when either population size or range size was included as a weight (See Tables 1 and 2).

### Mammals Article 17

We did not find any evidence that the probability that mammal populations were reported as increasing varies with either the diversity of conservation measures or threats, over either the short term or the long-term time span (Table 3). Similarly, we found no evidence for a relationship between the magnitude of change in population size and either diversity of threats or conservation measures reported (Table 3). The impact of time span on these response variables varied, with considerable uncertainty (Fig S6), and the clearest relationship being that although the change in population size of these selected mammals is positive (>0), it is less positive over greater time spans (between 25 and 30 years, Fig S6 D).

**Table 3.** Model comparison for analyses of trends in mammal population size. P(increasing) = probability that population size was reported to be increasing, Mag(increase) = Magnitude of change in population size reported. Best model (lowest AIC) given in bold. If the set of best models included the null model, we follow a conservative approach and consider the null model as best.

| Response | Time Period | Formula | Weights | AIC | deltaAIC | Relative LL | AICweights |
| --- | --- | --- | --- | --- | --- | --- | --- |
| **P(increasing)** | **short** | **time_span + (1 \| speciesname)** | **none** | **78.35** | **0.00** | **1.00** | **0.31** |
| P(increasing) | short | measures + time_span + (1 \| speciesname) | none | 79.22 | 0.86 | 0.65 | 0.20 |
| P(increasing) | short | threats * measures + time_span + (1 \| speciesname) | none | 79.25 | 0.90 | 0.64 | 0.20 |
| P(increasing) | short | threats + measures + time_span + (1 \| speciesname) | none | 79.74 | 1.39 | 0.50 | 0.16 |
| P(increasing) | short | threats + time_span + (1 \| speciesname) | none | 80.13 | 1.77 | 0.41 | 0.13 |
| **P(increasing)** | **long** | **time_span + (1 \| speciesname)** | **none** | **34.79** | **0.00** | **1.00** | **0.48** |
| P(increasing) | long | measures + time_span + (1 \| speciesname) | none | 36.41 | 1.63 | 0.44 | 0.21 |
| P(increasing) | long | threats + time_span + (1 \| speciesname) | none | 36.77 | 1.98 | 0.37 | 0.18 |
| P(increasing) | long | threats + measures + time_span + (1 \| speciesname) | none | 38.16 | 3.38 | 0.18 | 0.09 |
| P(increasing) | long | threats * measures + time_span + (1 \| speciesname) | none | 40.10 | 5.31 | 0.07 | 0.03 |
| Mag(increase) | short | threats + time_span + (1 \| speciesname) | none | 99.39 | 0.00 | 1.00 | 0.53 |
| **Mag(increase)** | **short** | **time_span + (1 \| speciesname)** | **none** | **100.84** | **1.45** | **0.48** | **0.26** |
| Mag(increase) | short | threats + measures + time_span + (1 \| speciesname) | none | 102.64 | 3.24 | 0.20 | 0.10 |
| Mag(increase) | short | measures + time_span + (1 \| speciesname) | none | 103.29 | 3.90 | 0.14 | 0.07 |
| Mag(increase) | short | threats * measures + time_span + (1 \| speciesname) | none | 104.58 | 5.19 | 0.07 | 0.04 |
| **Mag(increase)** | **long** | **time_span + (1 \| speciesname)** | **none** | **49.27** | **0.00** | **1.00** | **0.42** |
| Mag(increase) | long | measures + time_span + (1 \| speciesname) | none | 50.32 | 1.06 | 0.59 | 0.25 |
| Mag(increase) | long | threats + time_span + (1 \| speciesname) | none | 50.90 | 1.64 | 0.44 | 0.19 |
| Mag(increase) | long | threats * measures + time_span + (1 \| speciesname) | none | 52.17 | 2.90 | 0.23 | 0.10 |
| Mag(increase) | long | threats + measures + time_span + (1 \| speciesname) | none | 54.13 | 4.86 | 0.09 | 0.04 |

We were not adequately able to test which variables influenced the probability that the range of the selected mammals was reported as expanding (versus shrinking), over either the short term or long term, due to a very small number of data points on range contractions. The uncertainty around the modelled relationships was very high for the short term data (Fig S7). We were not adequately able to test these relationships with the corresponding long term data, as only four (of 54) data points gave information on range decreases, and model coefficients were not reliable (Fig S8). We could not adequately test for a relationship between either conservation measures or threats and the magnitude of mammal range expansion over the short term and found no evidence for a relationship between these variables over the long term (Table 4).

**Table 4.**
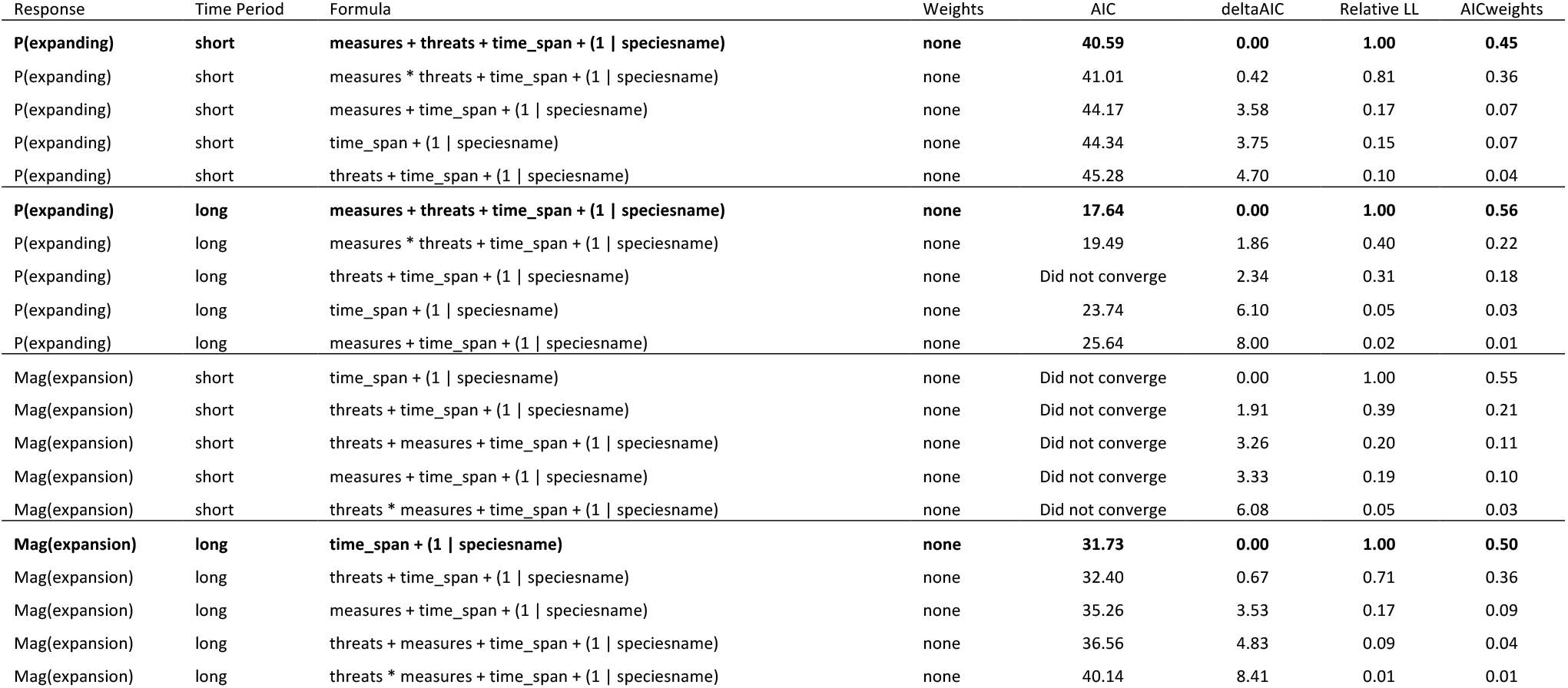
Model comparison for analyses of trends in mammal range. P(expanding) = probability that population range was reported as expanding, Mag(expansion) = Magnitude of change in range reported. Best model (lowest AIC) given in bold. If the set of best models included the null model, we follow a conservative approach and consider the null model as best.

### Mammal abundance trends in the Living Planet Index Database

Two models were supported with similar strength following selection based on AIC values (Table 5, Fig 3). Both these models suggest that the relative change in population size of recovering mammals is greater where those populations are managed, and lower where the population is utilised or known to be impacted by threats (see Fig 3 C to E). Where the managed, utilised or threatened status was unknown, the impact was intermediate between “yes” and “no”. The relative change in population size is slightly lower for larger mammals, although this relationship was only supported by one of the two best models (Fig 3 B). There is also an increase in the relative change in population size with longer time series (Fig 3 A). Neither of the best models included protection status, suggesting that this is not strongly influencing changes in the population size of the focal recovering mammal species. The estimates and 95% CI for estimates of the average of the two best models is given in Fig S9 and shows the same pattern of results.

**Table 5.**
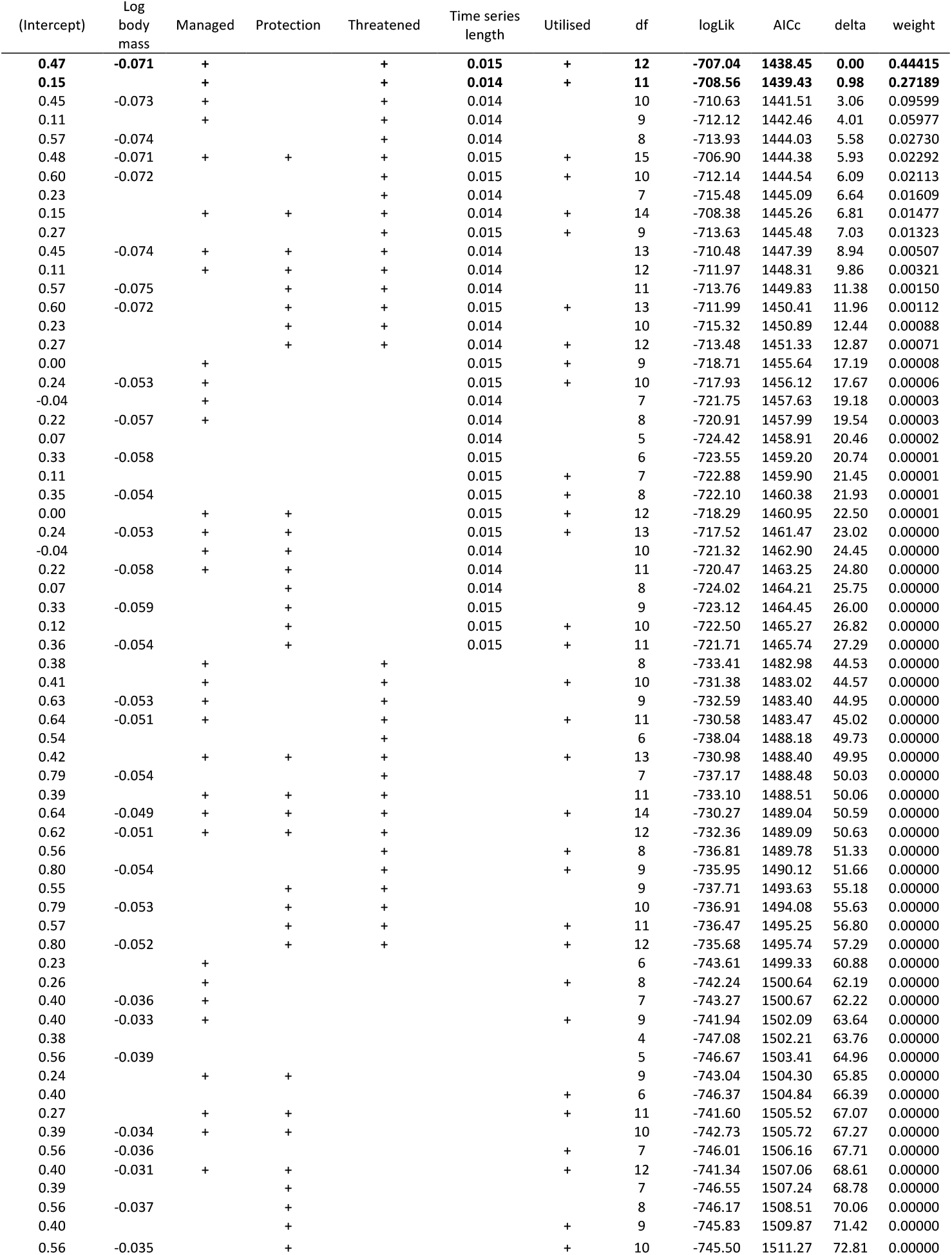
Model comparison for total lambda for mammal species. Best model (lowest AIC) given in bold, multiple models are highlighted where the difference in AIC was <2.

**Fig 3.**
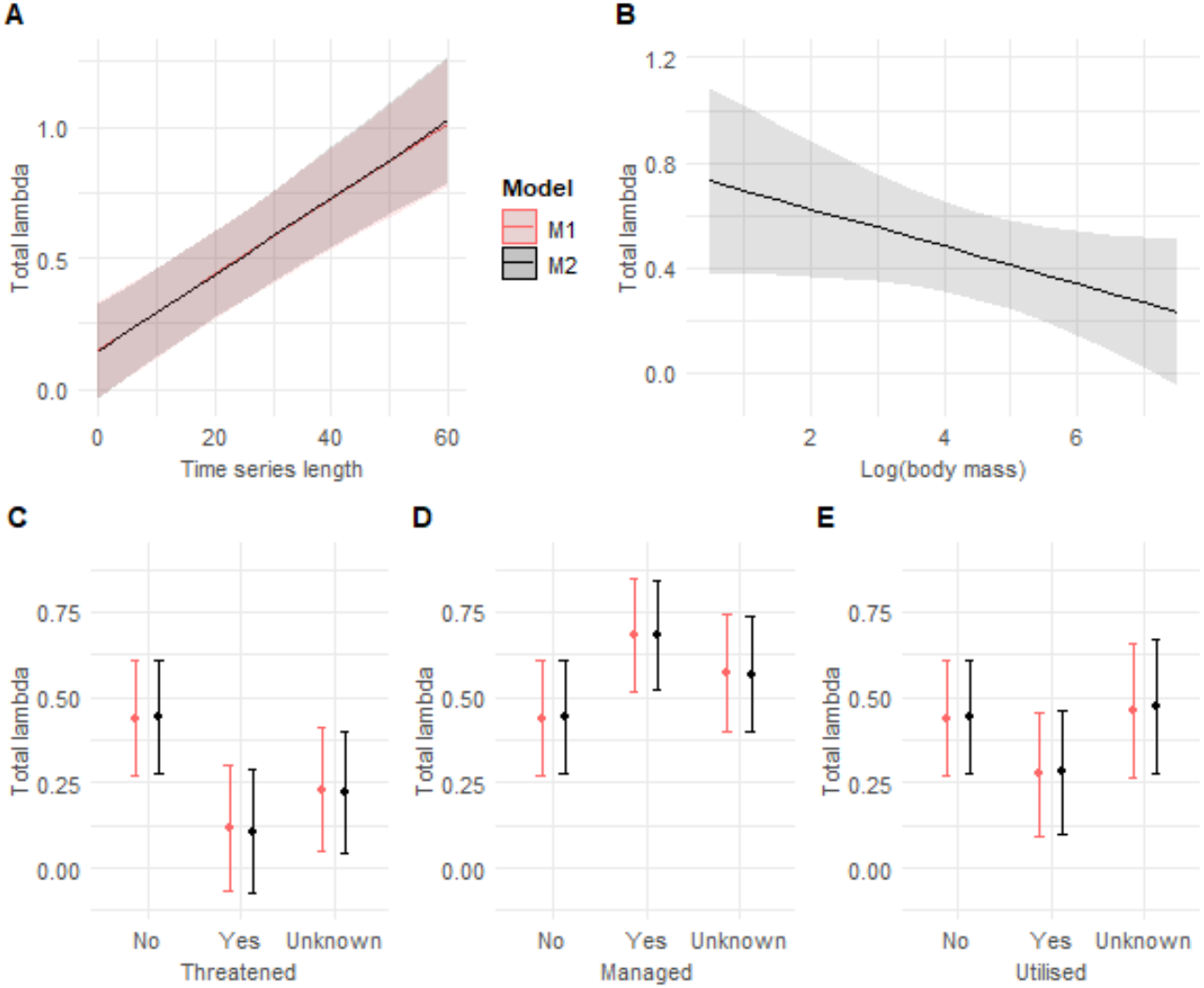
The relative abundance of selected mammal species A) increases with the length of the data time series, B) decreases slightly with increasing body mass and varies with the presence of C) threats, D) targeted managed and C) use or offtake of the species. Plots show predicted values and 95% CI for the predictor variables in the two best models with similar support. Panels A, C, D, E show M1 (best model, see Table 5) in red and the second model with equal support, M2, in black. Panel B shows only the predicted values from M2 as it was only this model that contained the variable for body mass.

## Discussion

As we move into the UN Decade on Restoration, it is especially important to look in detail at cases of species recovery, so that we can understand the drivers behind these changes and use this information to identify what is crucial for further success and what may be limiting existing recoveries.

Overall, our results show that even for recovering species, increases in population size and range are lower where there are more threats in place, and that these species have clearly benefited from conservation measures. Over the longer time spans reported by EU Member States (generally 1980 to 2018), the breeding birds we selected as currently “recovering” are doing less well (in terms of both range and population size) when they are impacted by a greater diversity of threats. Although we emphasise the results of the simplest, or most parsimonious model, there was also some indication that the negative impact of threats may be mitigated by a greater diversity of conservation measures. These interactions suggest that the negative effect of a greater diversity of threats was more pronounced when a small number of conservation measures were reported and was less marked when more conservation measures were in place. Although we were not able to draw conclusive results from the data reported under Article 17, these results are further reinforced for mammals by the analysis of data from the LPD (which starts in 1960 and is therefore more comparable to the long-term time period). We found that increases in abundance are less positive where there is direct offtake for human use or other known threats in place. The same mammal populations also show more positive increases in abundance where they are known to be managed. These results emphasise the importance of continuing to reduce threats (such as habitat loss, hunting, invasive species, climate change) even for those species that are known to be “coming back” across Europe, and continuing to support the conservation measures that are in place.

Although the available data did not allow as thorough testing of short-term relationships, we did not see evidence for relationships between the diversity of threats or conservation measures and the majority of population and range variables calculated from the EU reported data. The short-term time span in the EU data is generally 2007 to 2018, so this difference could be explained by the generation lengths of the species selected; any changes in measures or threats are unlikely to have caused large changes in population increase or range expansion if only affecting one or two generations. Indeed, IUCN Red List assessments of extinction risk using abundance trends (criterion A) would consider trends over 3 generation lengths. Many of the bird species in this exercise have long life spans; the average generation length of the 23 species included here is 10.6 years, with a standard deviation of 4.28, using values from (Bird et al. 2020). We would therefore only expect threats or conservation measures having a high, direct impact on mortality to drive a noticeable change over the c. 12 years covered by the short-term time span. Similarly, the average generation length of the mammal species included here is 9.6 years (standard deviation of 4.4) (Pacifici et al. 2013). Ideally in the future it would also be useful to be able to better explore the effect of different types of conservation measures and threats, as some (e.g. hunting bans) may act over shorter timescales than others (establishing protected areas or initiating ecosystem restoration). Our summary of the number of countries reporting different threats and measures (Fig 1) shows that some types of measures are much more widespread than others, and it would be important to look further and how the timescale over which a measure has impact relates to the scale at which it is applied.

Given the weak support for some interactions between threats and measures in the breeding bird data reported under Article 12, future research (ideally with more detailed data for more populations) would also benefit from examining where threats or conservation measures may act synergistically. We were also not able to examine the difference between species that are suffering from a greater diversity of threats versus a greater intensity of a reduced range of threats. Future analyses drawing on data that captures the type, timing, severity and scope of threats, such as that stored through the threat classification scheme of the IUCN Red List (IUCN 2022), may be able to address these issues; an important step to inform strategies to roll out threat mitigation or conservation measures.

A key conservation measure of interest is that of protected areas. Protected areas can reduce human threats and benefit biodiversity, although the impact varies with geographic region and ecosystem type (e.g. Butchart et al. 2012; Geldmann et al. 2013, 2019; Gray et al. 2016) and depends on adequate resourcing (Geldmann et al. 2018). We were only explicitly able to look at the importance of protection for mammals (using the LPD), and we note that there was no support for a relationship between protection status and relative increase in the abundance of the selected mammal populations. However, our results are consistent with recent research showing that the extent of protection and human footprint has very little influence on the distribution of 13 European carnivores and ungulates (Cretois et al. 2021), 11 of which are also included in the LPI subset we analysed. Indeed, the European protected area network is small compared to the range size required by many large mammals, and is also made up of cultural landscapes with multiple uses allowed and preserved, such that large mammals thrive equally well outside the reserves (Linnell et al. 2015). Our results do suggest that active management (which includes conservation measures such as reintroduction and protection of the species itself rather than the area it is in) was positively related to mammal abundance, indicating that targeted management is more important for the recovery of European mammals than the protected area network. Although we were not able to explicitly assess the importance of the protected area system for our selected recovering bird species, there is strong evidence that the Birds Directive, Natura 2000 and wider protected area networks are considered to be a key driver of improvements in European bird species (Donald et al. 2007; Tucker et al. 2019).

Our analyses highlight that even with coordinated reporting across a well-resourced set of countries, there are still substantial limitations due to data availability. The data we have suffers from inaccuracies and variation due to differences in the level of capacity that Member States have to deliver the data (countries vary considerably in the funding available and the availability of skilled amateur volunteers (Schmidt et al. 2021). Of the species reported on under the Birds Directive, a third of the Article 12 data (country-level reporting on bird population sizes and trends) are based on complete surveys, one fifth are based on expert opinion, and around 40% have partial, limited or no data available. Crucially, here our taxonomic breadth was limited to only birds and mammals. The critical need to keep improving data availability is emphasised throughout the literature on wildlife recovery across Europe; with much better insights needed into marine, freshwater and underground ecosystems (Carver et al. 2021). To fully understand the drivers of decline and recovery we also need to improve the coordination of data on threats and measures across Europe, such as hunting, vehicle collisions and damage/compensation payments (Linnell et al. 2020). These data need to be robustly monitored over suitable time-periods and using well-established shared frameworks describing threats and conservation measures. There has been some progress towards this with Article 12-like reporting processes now been adopted by the Africa-Eurasian Waterbird Agreement (AEWA) and trialled by Bern Convention, i.e. 6-yearly updates. There are examples of some initiatives that are helping achieve these aims. EuropaBON proposed solutions to include funding, capacity building, technology, coordination and standardisation (Moersberger et al 2022). Embedding monitoring within policy frameworks can help e.g. aquatic monitoring has improved under the auspices of the Water Framework Directive, which helped with standardisation (Moersberger et al 2022).

The species featured here have been recovering from a wide range of actions, including translocation/reintroductions, legal-protections, site-management, educational programs, changes as people move back into cities. Tucker et al (2019) identified a range of key drivers of success across 53 case studies of species with genuine improvements from the Habitats and Bird Directives. Across these case studies they also highlighted important high-level drivers, including strong, coherent governance, effective institutions and committed individuals to support implementation, increased habitat and species protection and a range of other important recommendations. Transboundary management is also important, especially for migratory species (Linnell et al, 2020). Protection of breeding, passage and wintering sites has been fundamental to the recovery of some migratory bird species (e.g. Deinet et al. 2013); ensuring such effective protection for longer-distance migrants can be more complex, especially when their movements take them outside Europe.

Here, our results provide evidence that the *diversity* of current threats is a factor in inhibiting recoveries. Reducing threat diversity may therefore be a priority. However, as new threats continue to emerge, this may be complex, particularly as action can be hard to instigate until negative impacts have been recorded. How threat richness interacts with the type, severity and scope of those threats also remains to be quantified.

The small number of selected recovering species we assess here sit within the context of global biodiversity declines and ongoing extinctions. Understanding the processes that drive and limit recoveries is critical if we wish to ensure that a greater range of species and ecosystems can recover to historically healthy levels and enable us to achieve the aims of the UN Decade on Restoration. Here, we explored how existing threats and conservation measures influence the likelihood and magnitude of recoveries in selected, recovering European birds and mammals. Our results demonstrate that even considering these recovering species, threat diversity can continue to limit longer-term increases in population size and range extent, and perhaps this can be mitigated using a range of measures/management. This highlights opportunities to further maximise the gains these species have begun to make, but also underlines the need to remain vigilant to ongoing and emergent threats that may limit potential recoveries.

## Acknowledgements

This work was funded in part by Rewilding Europe. L.M. was additionally funded by WWF UK. R.F. was funded by Research England. The authors are extremely grateful to the many data providers and data inputters to the Living Planet Database who have made this analysis possible.

## Author Contributions

RF, CG, LM, SD, SL: conceived of the original idea, curated, and analysed the data and prepared the original draft. KSG: collected additional abundance data. CB, HP and KSG: compiled additional research on species used in the analysis. All authors contributed to subsequent writing, reviewing and editing of the manuscript.

## Supplementary figures and tables

**Table S1.**
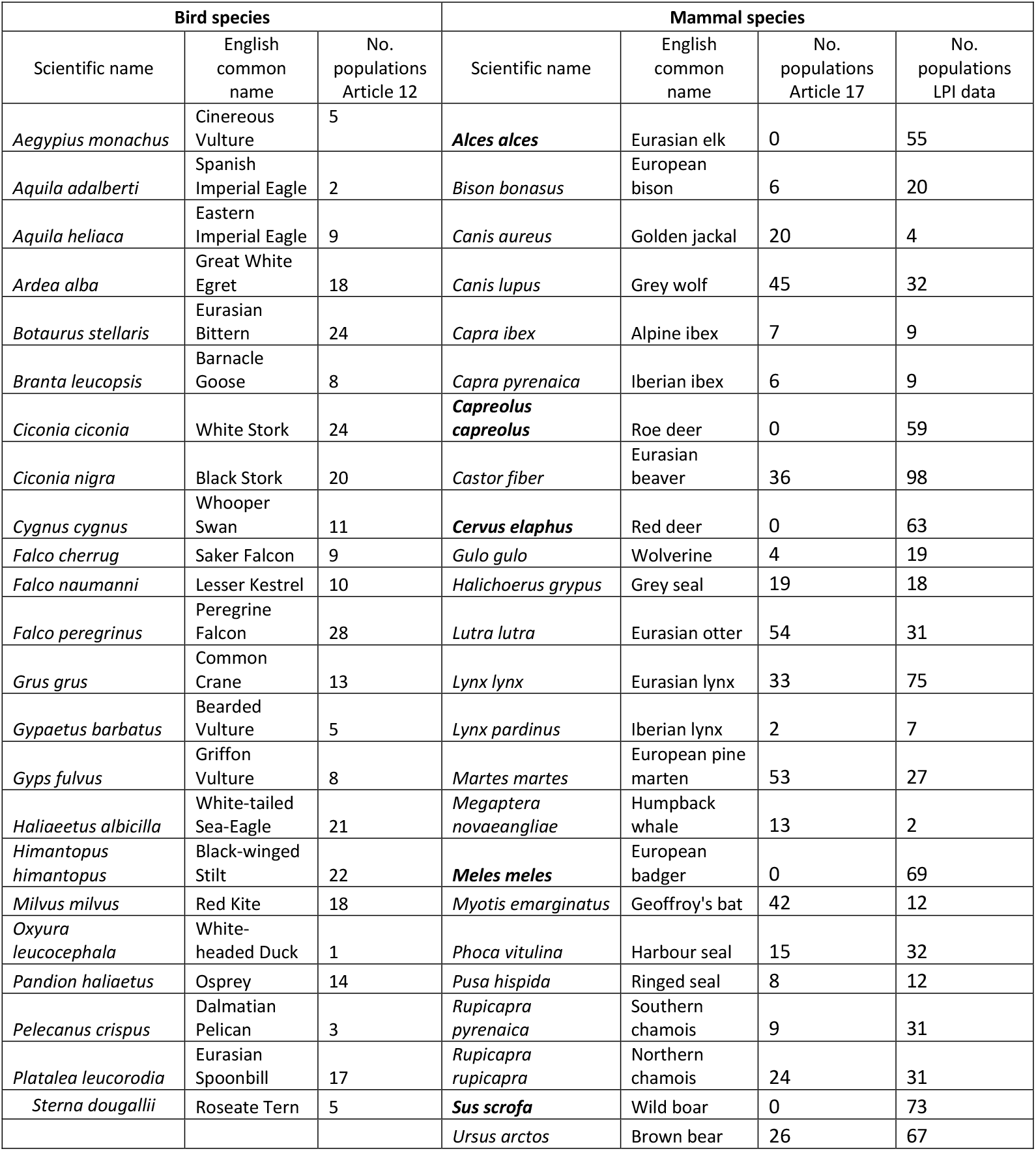
Bird and mammal species selected for the analysis. LPI data was available for all mammals listed; Article 17 data was not for the five species highlighted in bold. Article 12 data on was available for all species listed; number of populations refers to the reported data on breeding season only (wintering season was excluded).

**Table S2.** Size of datasets used for analysis of changes in selected bird (top) and mammal (bottom) species. For assessments of the probability and magnitude of abundance increase (P(increasing and Mag(increase) and the probability and magnitude of Range expansion (P(expanding) and Mag(expansion), the number of populations for both short-term and long-term analysis is shown.

| Birds | Analysis | Period |  |
| --- | --- | --- | --- |
|  |  | Short term (n) | Long-term (n) |
| **Abundance** | *P*(increasing) | 156 | 198 |
|  | *Mag*(increase) | 153 | 136 |
| **Range-change** | *P*(expanding) | 141 | 163 |
|  | *Mag*(expansion) | 163 | 169 |

| Mammals | Analysis | Period |  |
| --- | --- | --- | --- |
|  |  | Short term (n) | Long-term (n) |
| **Abundance** | *P*(increasing) | 137 | 70 |
|  | *Mag*(increase) | 39 | 24 |
| **Range-change** | *P*(expanding) | 93 | 54 |
|  | *Mag*(expansion) | 16 | 16 |

**Table S3.**
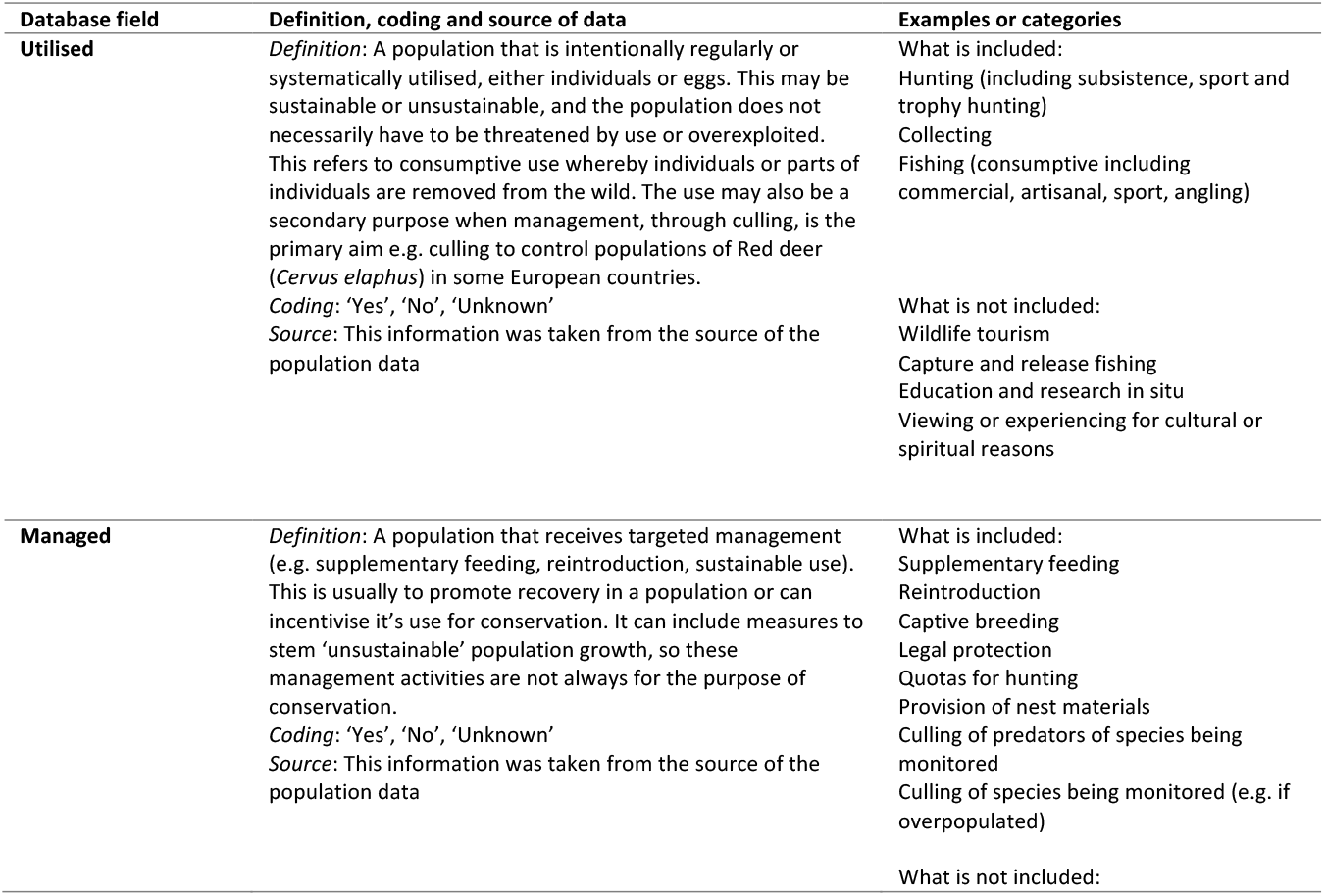
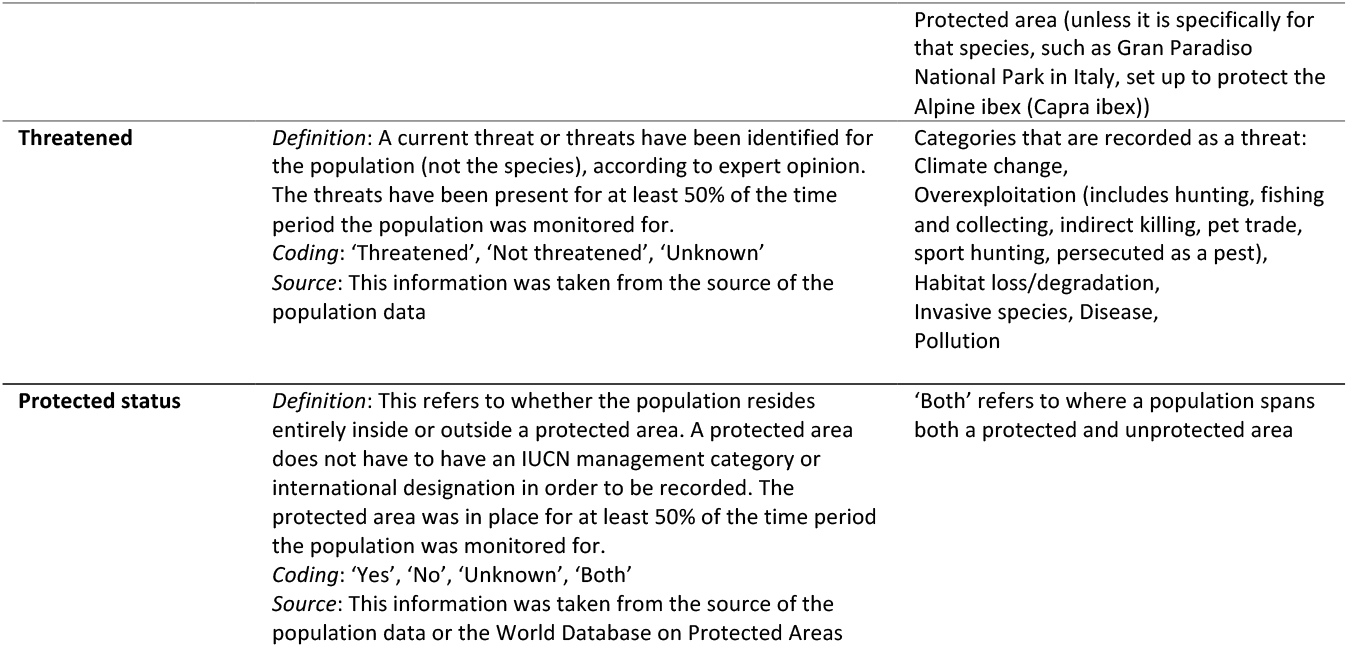
Ancillary data fields used for modelling mammal population data from the LPI Database.

**Table S4.**
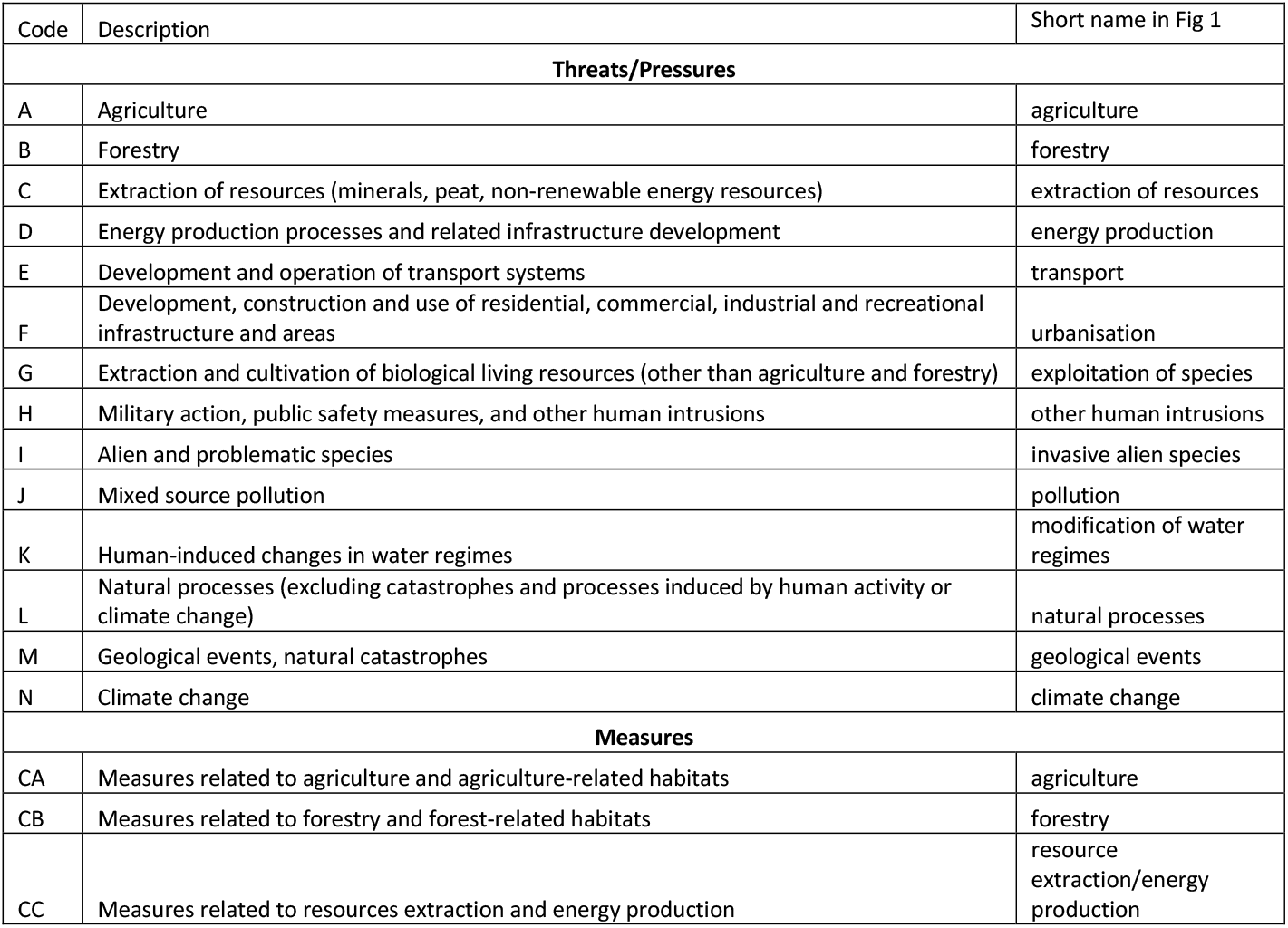
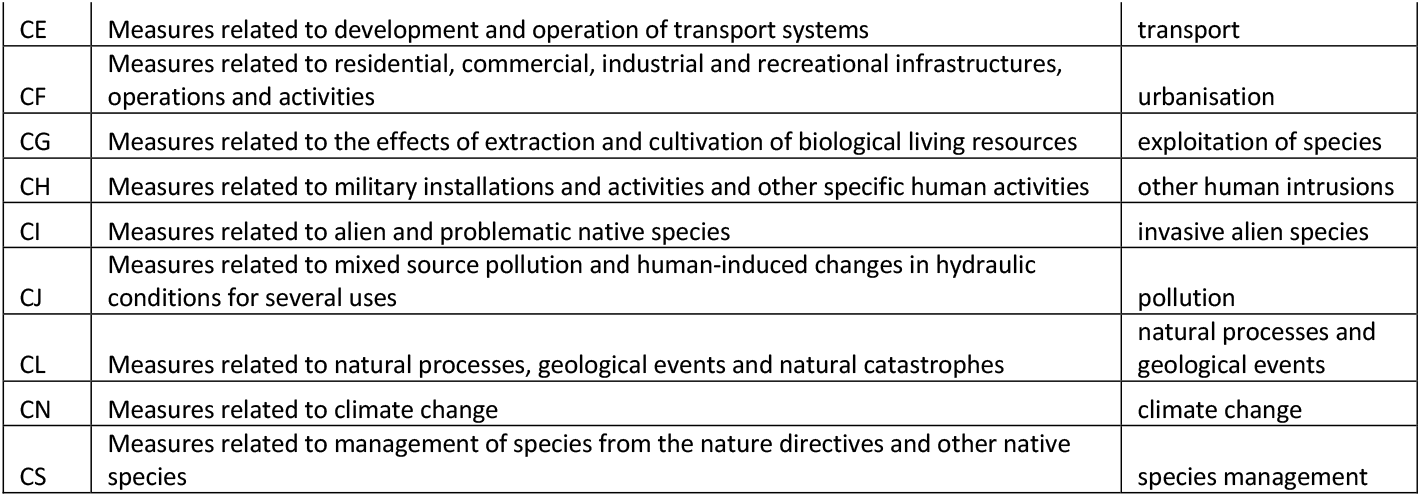
Threat/pressure and measure codes used by EU for data reported under Article 12 of Bird Directive and Article 17 of Habitats Directive; our metrics for diversity of threats and conservation measures equal the total number of these codes reported for a species. Our use of short names follows (Röschel et al. 2020).

| Code | Description | Short name in Fig 1 |
| --- | --- | --- |
| **Threats/Pressures** |  |  |
| A | Agriculture | agriculture |
| B | Forestry | forestry |
| C | Extraction of resources (minerals, peat, non-renewable energy resources) | extraction of resources |
| D | Energy production processes and related infrastructure development | energy production |
| E | Development and operation of transport systems | transport |
| F | Development, construction and use of residential, commercial, industrial and recreational infrastructure and areas | urbanisation |
| G | Extraction and cultivation of biological living resources (other than agriculture and forestry) | exploitation of species |
| H | Military action, public safety measures, and other human intrusions | other human intrusions |
| I | Alien and problematic species | invasive alien species |
| J | Mixed source pollution | pollution |
| K | Human-induced changes in water regimes | modification of water regimes |
| L | Natural processes (excluding catastrophes and processes induced by human activity or climate change) | natural processes |
| M | Geological events, natural catastrophes | geological events |
| N | Climate change | climate change |
| **Measures** |  |  |
| CA | Measures related to agriculture and agriculture-related habitats | agriculture |
| CB | Measures related to forestry and forest-related habitats | forestry |
| CC | Measures related to resources extraction and energy production | resource extraction/energy production |
| CE | Measures related to development and operation of transport systems | transport |
| CF | Measures related to residential, commercial, industrial and recreational infrastructures, operations and activities | urbanisation |
| CG | Measures related to the effects of extraction and cultivation of biological living resources | exploitation of species |
| CH | Measures related to military installations and activities and other specific human activities | other human intrusions |
| CI | Measures related to alien and problematic native species | invasive alien species |
| CJ | Measures related to mixed source pollution and human-induced changes in hydraulic conditions for several uses | pollution |
| CL | Measures related to natural processes, geological events and natural catastrophes | natural processes and geological events |
| CN | Measures related to climate change | climate change |
| CS | Measures related to management of species from the nature directives and other native species | species management |

**Fig S1.**
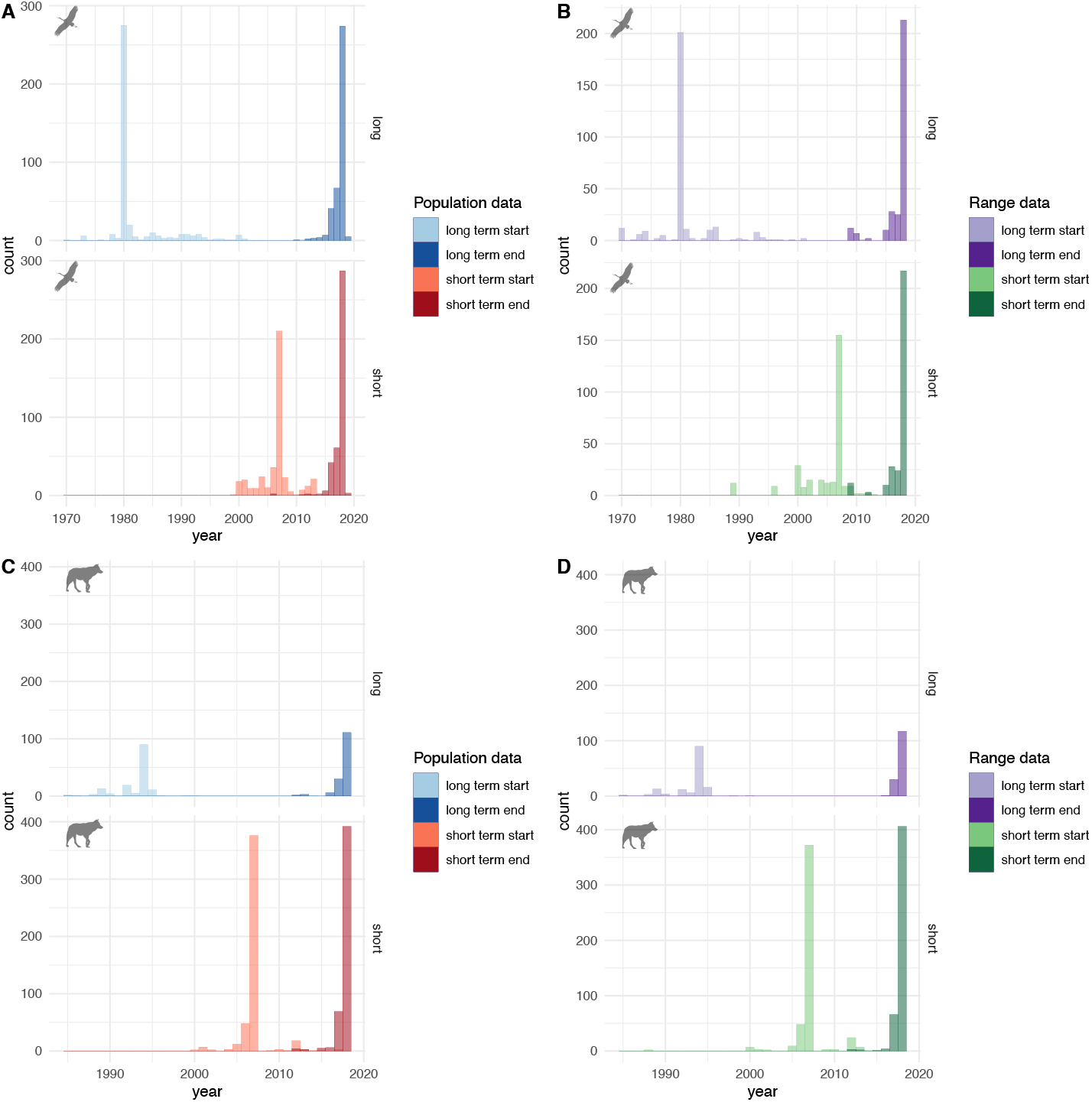
Distribution of start and end years for the data was reported under Article 12 for bird species’ population size (A) and range (B) and under Article 17 for mammal species’ population size (C) and range (D). Upper and lower plots in each panel show distributions for short and long term reporting periods.

**Fig S2.**
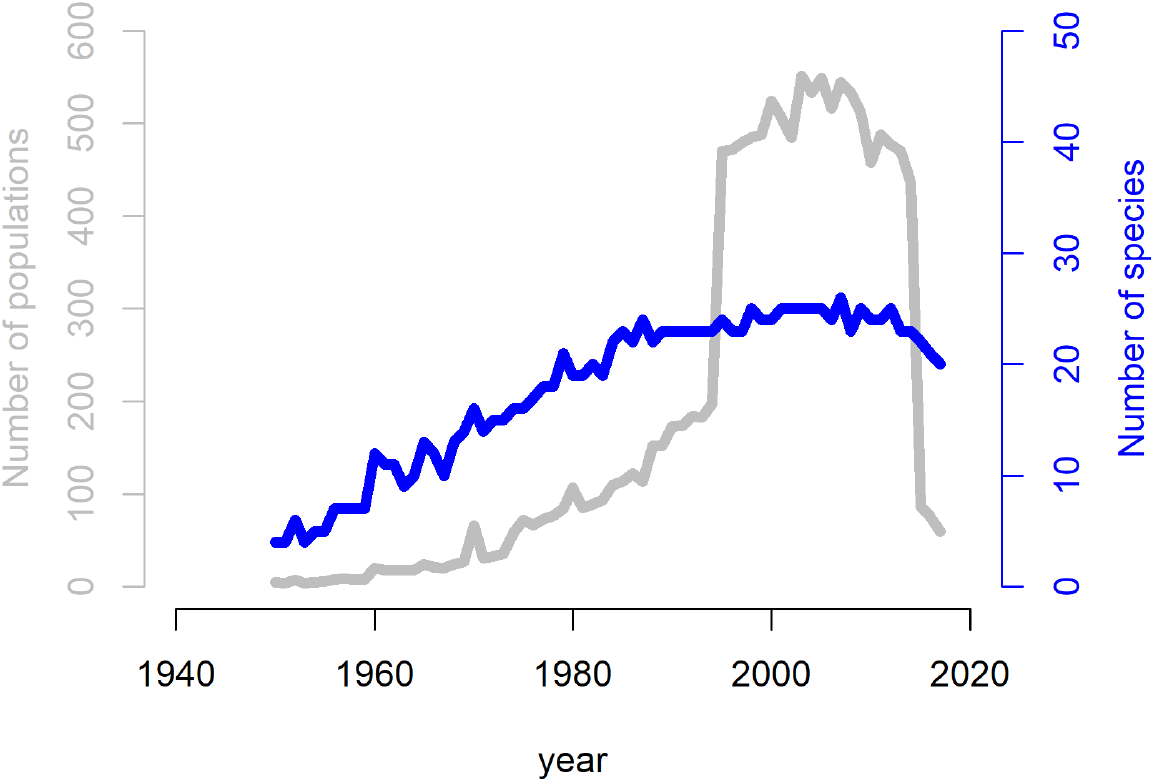
Total number of populations (grey) and species (blue) for which the LPD contains abundance data for each year between 1950 and 2016, for the European mammals considered in this study

**Fig S3.**
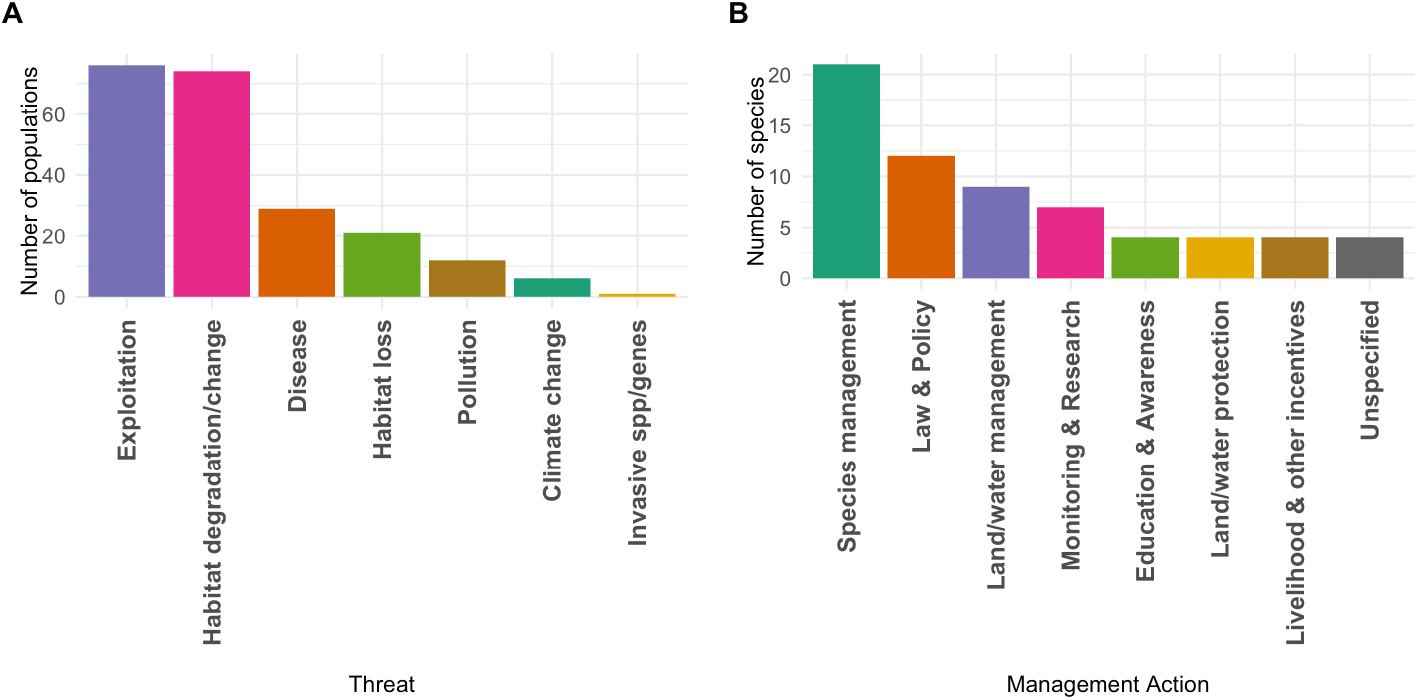
A) Frequency of identified threats for mammal populations, and (B) identified population management information within the Living Planet database. Categories were assigned using the IUCN Red List Threat classification scheme or IUCN Red List Conservation action classification (after Jellesmark et al 2022).

**Fig S4.**
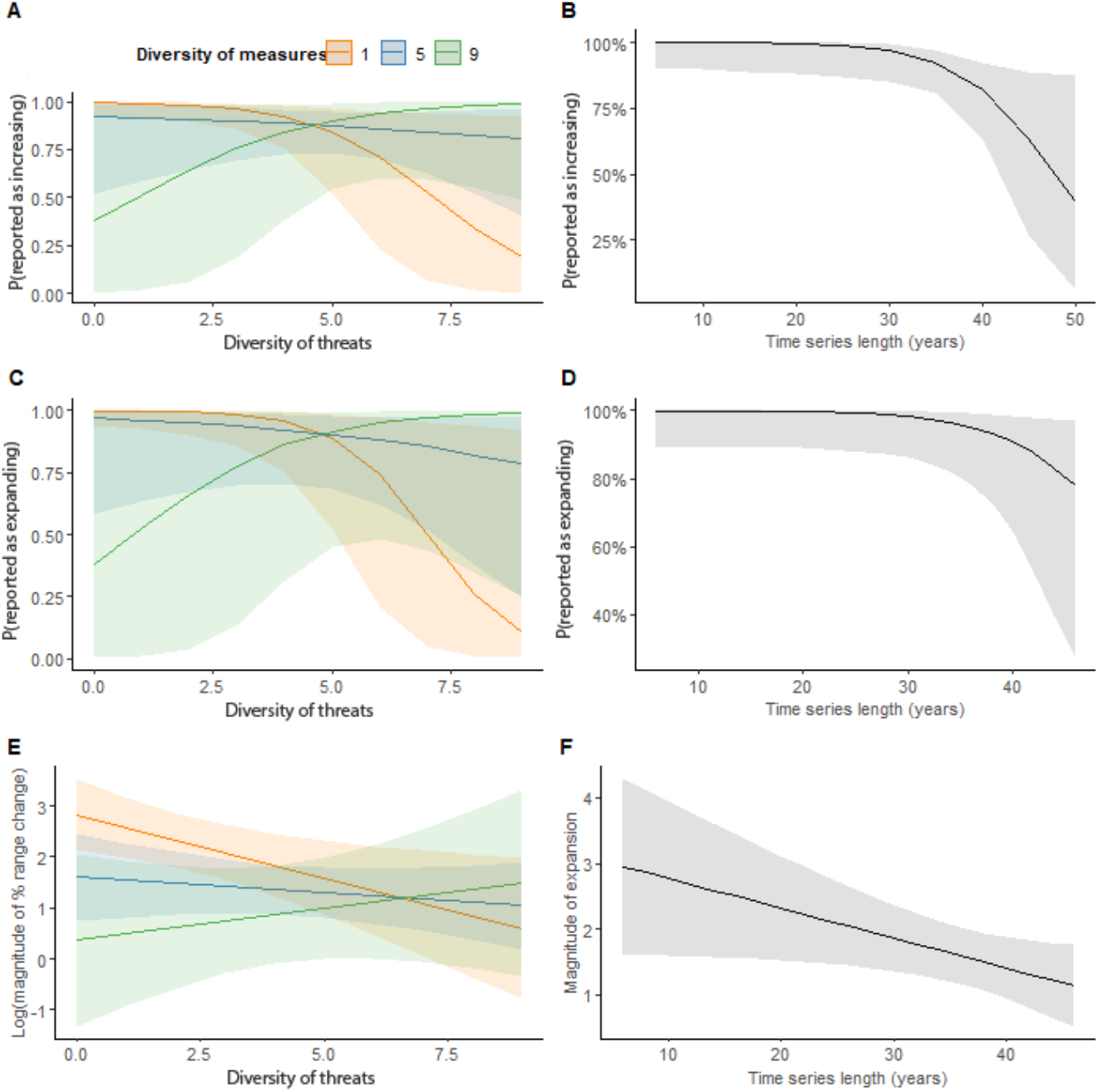
There is some evidence that the interaction between the diversity of threats and diversity of conservation measures influences A) the probability that selected breeding bird populations are increasing over the long term, C) the probability that these species ranges are reported as expanding over the long term and E) the magnitude of the range expansion over the long term. As for the simpler model only including threats (Fig. 2), there is increasing uncertainty around the impact of the time span of the data when it is greater than 40 years for B) the probability that selected breeding bird populations are increasing over the long term and D) the probability that these species ranges are reported as expanding. Again, similar to the simpler model only including threats (Fig. 2), there is evidence that the F) magnitude of expansion over the long term decreases as time span increases.

**Fig S5.**
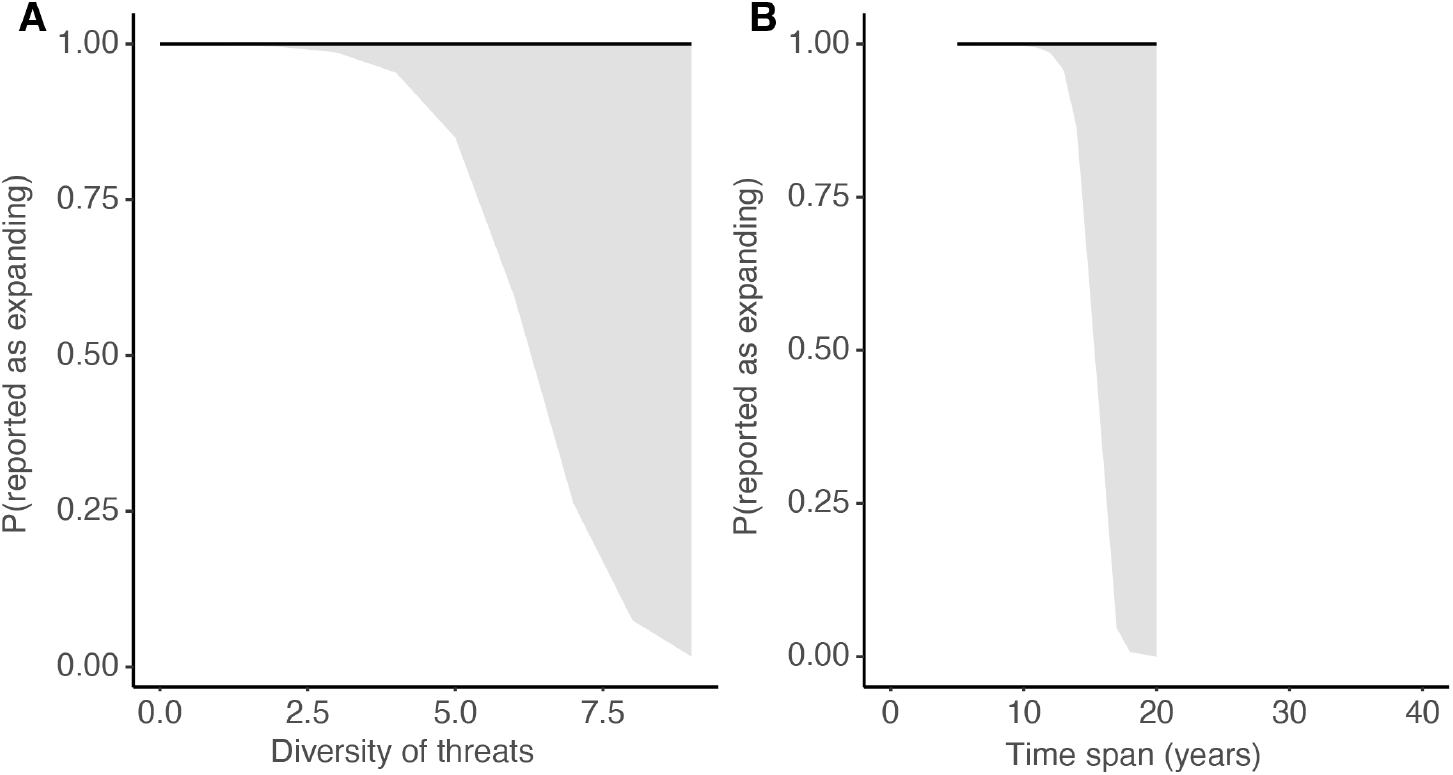
The probability that the selected breeding bird ranges are expanding (over the short term) does not have a clear relationship with either A) the diversity of threats reported or B) the time span of the data.

**Fig S6.**
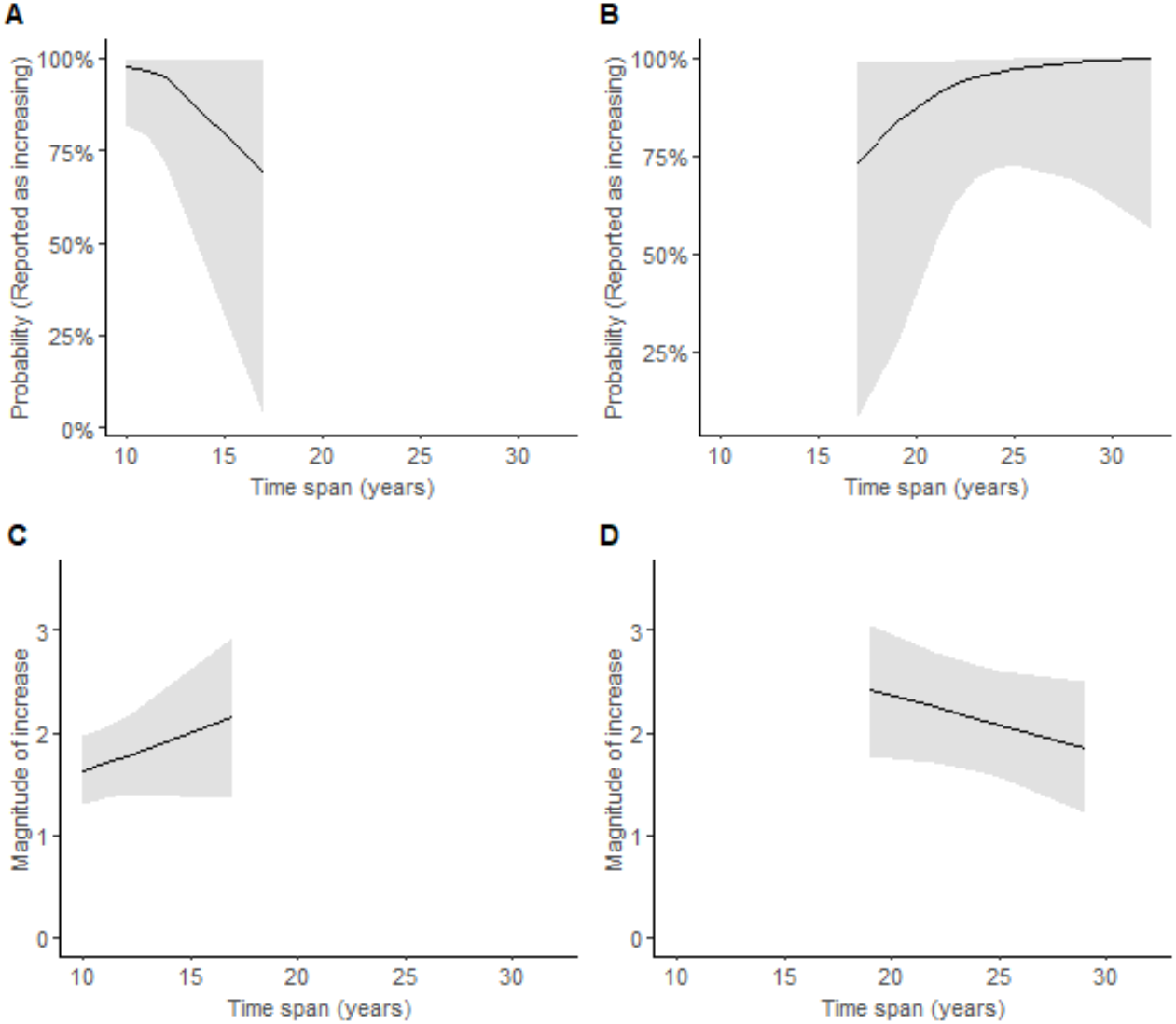
Modelled relationship between the probability that the population of selected mammals was reported as increasing over the A) short term and B) long term, and the magnitude of the mammal population increase over the C) short term and D) long term.

**Fig S7.**
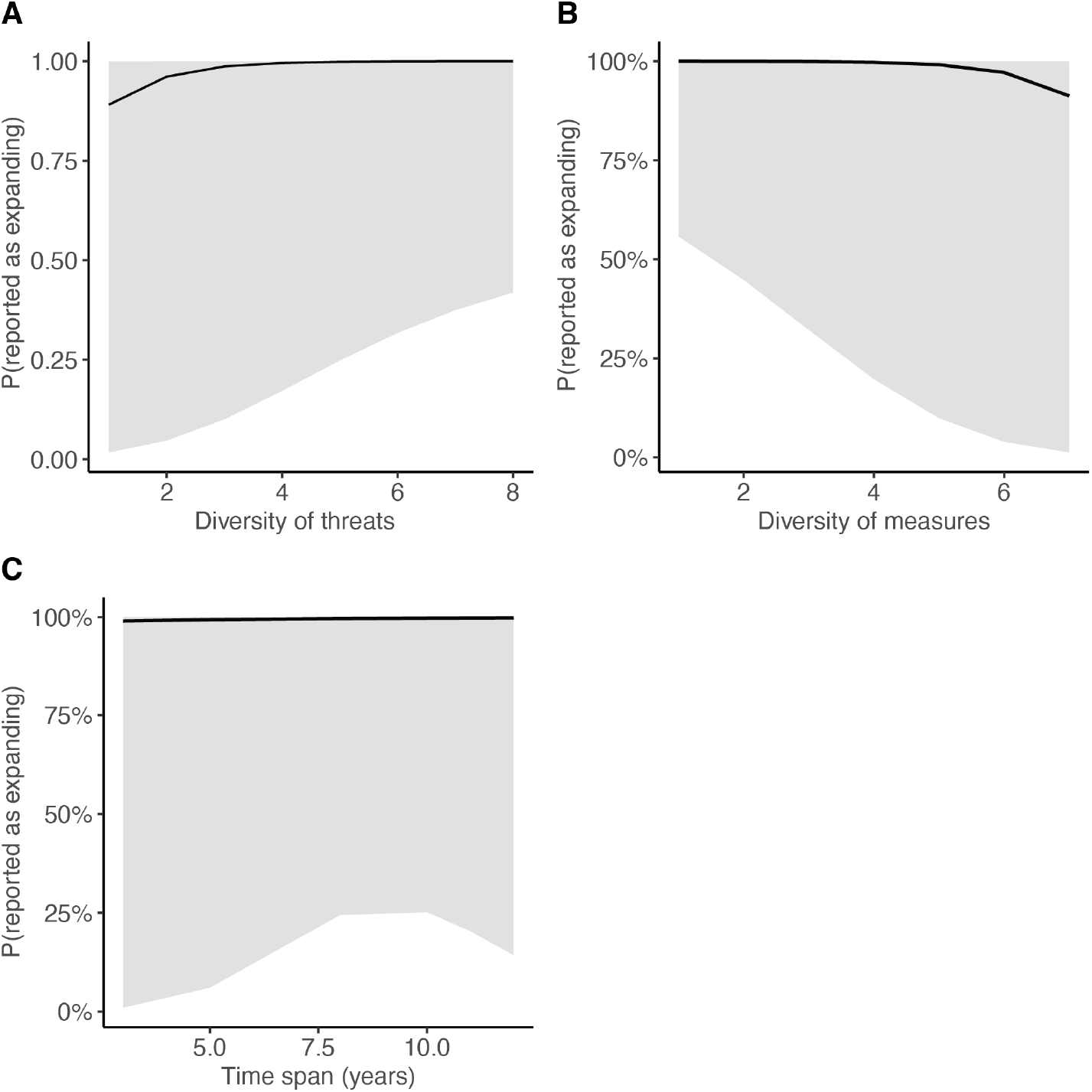
For our selected mammal species, we found high uncertainty around the modelled relationships between the probability of an expansion in range over the short term and the diversity of threats (A), diversity of conservation measures (B) and time span (C). Very few data points gave information on a decrease in range, hence we do not consider the data to be adequate to properly test for these relationships.

**Fig S8.**
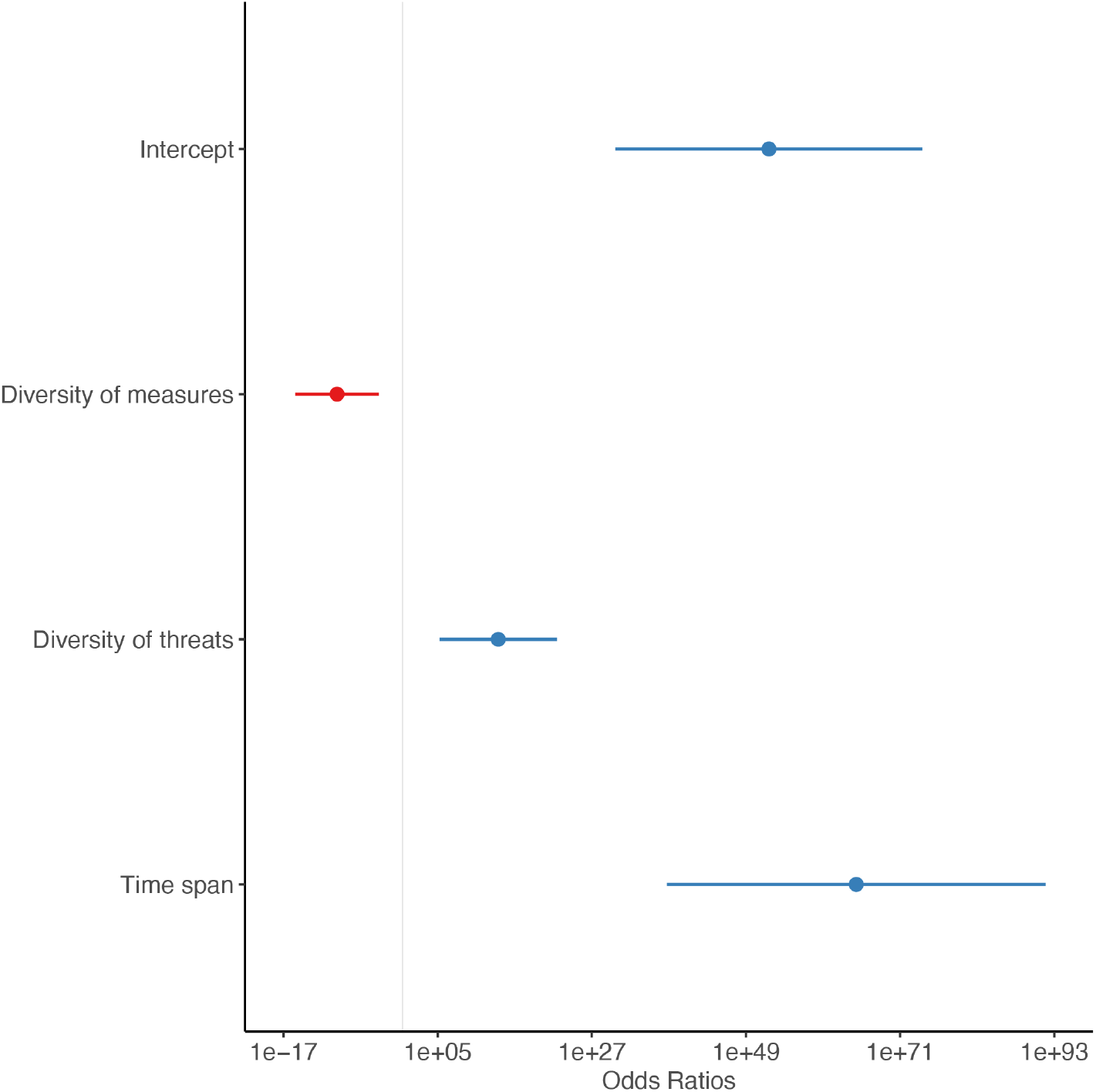
Coefficient plot for the model of Article 17 data for selected mammal species, showing particularly high odds ratios for the relationship between diversity of threats, diversity of measures, time span and the response variable P(expanding) over the long term. This model was run on data with only 4 (out of 54) data points for a decrease in range, hence we do not consider the data to be adequate to properly test for these relationships.

**Fig S9.**
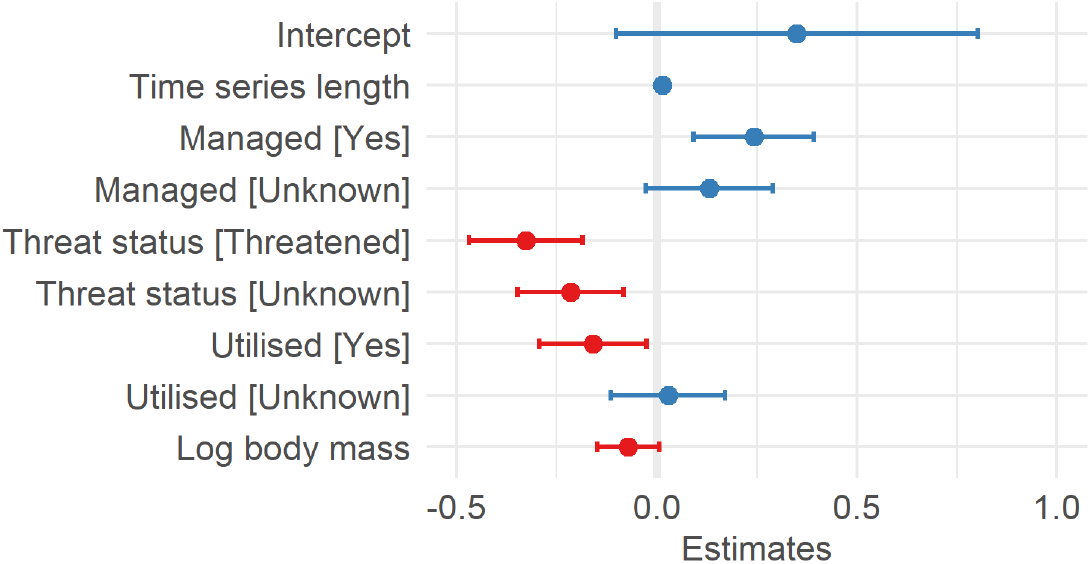
Estimates from the average of the two most strongly supported models testing for relationships between the predictor variables and the relative increase in abundance of selected mammal species.

## Notes

### Competing Interest Statement

The authors have declared no competing interest.

